# Proximity interactome of LC3B in normal growth conditions

**DOI:** 10.1101/2021.10.08.463639

**Authors:** Marie Nollet, Alexander Agrotis, Fanourios Michailidis, Arran David Dokal, Vinothini Rajeeve, Jemima Burden, Thomas D. Nightingale, Pedro Cutillas, Robin Ketteler, Stéphanie Kermorgant

**Affiliations:** Spatial Signalling Team, Barts Cancer Institute, Queen Mary University of London, John Vane Science Centre, Charterhouse Square, London EC1M 6BQ, United Kingdom; MRC Laboratory for Molecular Cell Biology, University College London, London WC1E 6BT, United Kingdom; Cell Signalling & Proteomics Group, Barts Cancer Institute, Queen Mary University of London, EC1M 6BQ London, United Kingdom; Centre for Microvascular Research, William Harvey Research Institute, Barts and the London School of Medicine and Dentistry, Queen Mary University of London, EC1M 6BQ London; MRC Protein Phosphorylation & Ubiquitylation Unit, School of Life Sciences, University of Dundee, DD1 5EH Dundee, United Kingdom; The Francis Crick Institute: 1 Midland Rd, NW1 1AT London

**Keywords:** autophagy, APEX2, LC3B, proximity proteomics

## Abstract

LC3 (Light Chain 3) is a key player of autophagy, a major stress-responsive proteolysis pathway promoting cellular homeostasis. It coordinates the formation and maturation of autophagosomes and recruits cargo to be further degraded upon autophagosome-lysosome fusion. To orchestrate its functions, LC3 binds to multiple proteins from the autophagosomes’ inner and outer membranes, but the full extent of these interactions is not known. Moreover, LC3 has been increasingly reported in other cellular locations than the autophagosome, with cellular outcome not fully understood and not all related to autophagy. Furthermore, novel functions of LC3 as well as autophagy can occur in cells growing in a normal medium thus in non-stressed conditions. A better knowledge of the molecule in proximity to LC3 in normal growth conditions will improve the understanding of LC3 function in autophagy and in other cell biology function. Using an APEX2 based proteomic approach, we have detected 407 proteins in proximity to the well-characterised LC3B isoform in non-stress conditions. These include known and novel LC3B proximity proteins, associated with various cell localisation and biological functions. Sixty-nine of these proteins contain a putative LIR (LC3 Interacting Region) including 41 not reported associated to autophagy. Several APEX2 hits were validated by co-immunoprecipitation and co-immunofluorescence. This study uncovers the LC3B global interactome and reveals novel LC3B interactors, irrespective of LC3B localisation and function. This knowledge could be exploited to better understand the role of LC3B in autophagy and non-autophagy cellular processes.

## Introduction

LC3B (Light Chain 3B), or MAP1LC3B (Microtubule-Associated Proteins 1A/1B light chain 3B), is the best-studied member of the seven protein-family LC3/GABARAP, and a homologue of yeast ATG8 (Autophagy Related 8) (1). The three isoforms LC3 A, B and C (hereafter LC3 when reporting on general functions or according to cited literature), are 17kDa proteins which localise on autophagosomes, the double membrane vesicles involved in the execution of macro-autophagy (hereafter, autophagy), a major intracellular proteolysis pathway (2). Autophagosome-lysosome fusion leads to the degradation of cytoplasmic material sequestered in the autophagosomes (3), releasing amino-acids for recycling and re-use by the cells (4). Autophagy is induced in response to stress, such as during infection by a pathogen, deprivation of nutrients or hypoxia, thereby promoting cell survival. However, if stress persists autophagy can lead to cell death. Cells can also display basal autophagy, allowing the clearance of long-lived proteins, damaged organelles and protein aggregates (5, 6). Autophagy is often upregulated in cancer cells (7, 8) and has been proposed to play a dual role in cancer: tumour suppressor in the early stages of tumorigenesis but pro-survival in advanced tumours (9).

LC3 is required for autophagosome formation and has been considered an autophagosome marker (10, 11). Autophagosome biogenesis (12) begins on the isolation membrane, or phagophore, with the assembly of a pre-initiation complex consisting of ULK1, FIP200, ATG101 and ATG13, which leads to activation of the VPS34/BECLIN1 PI3K Class III complex. Subsequent production of PI(3)P on the phagophore and binding of proteins such as WIPI2B leads to the recruitment of ATG16L1, which complexes with ATG5 and ATG12, forming a scaffold to attach LC3 to the forming autophagosome (13). For this to occur, LC3 is cleaved by ATG4B to generate LC3-I (cytosolic), capable of binding to ATG7, followed by transfer to ATG3. Finally, the ATG5-ATG12-ATG16L1 complex allows the covalent conjugation of LC3-I to phosphatidylethanolamine (PE), leading to LC3 lipidation, or the conversion of LC3-I to LC3-II (14, 15), which is required for autophagosome elongation and closure (16, 17).

LC3 is localised to the inner and outer membranes of the forming and mature autophagosome where it interacts with many proteins. This is often through the presence of a hydrophobic LC3 interacting region (LIR) (18–20). From the inner membrane, LC3 selectively recruits autophagic cargo into the autophagosome, binding the LIR of cargo receptors such as p62/SQSTM (21, 22), called selective autophagy. LC3 on the outer membrane of the autophagosome interacts with LIR-containing core autophagy machinery proteins such as ATG1/ULK1 (23), or proteins from the endolysosomal trafficking pathway such as FYVE and coiled-coil-domain-containing protein 1 (FYCO1) (21). This allows autophagosomes to traffic along microtubules and fuse with endocytic vesicles (autophagosome maturation), leading to the formation of degradative autolysosomes and autophagy (24, 25).

Non-conventional and non-autophagic functions of LC3 have also been increasingly reported (26). These can alter processes associated with membrane biology including the regulation of ER function (27, 28), secretion and exocytosis (29–34) and the non-canonical autophagy pathway LC3-Associated Phagocytosis (LAP) (35), which enables pathogen entry. LC3-positive vesicles may also play the role of signalling platforms (36–39). Non-autophagic functions of LC3 which do not appear related to membrane biology include the regulation of viral or bacterial replication (40, 41) and release (42) and pathogen control (43). The soluble form of LC3, LC3-I, is part of slowly diffusing high molecular weight complexes localised in the cytoplasm and the nucleus (44–46) but their function is poorly understood. Furthermore, LC3 was recently shown to conjugate to proteins (“LC3-ylation”) akin to ubiquitination (47), although only few candidate LC3-ylated proteins have been identified to date.

Despite the profound involvement of LC3 in cellular processes, autophagy-related or not, many of them remain poorly characterised. Consistent with this, the full repertoire of proteins interacting with LC3 is not known. Here we systematically defined the LC3B interactome by performing an enzymatic proximity tagging approach using an engineered ascorbate peroxidase, APEX2 (48, 49). We used HeLa cells stably transfected with an APEX2- and GFP-tagged LC3B construct (or the control GFP-LC3B) and cultured them in normal growth conditions. Our study has confirmed many proteins previously reported to be partners of LC3B and/or present in autophagosomes, including proteins with putative LIR. It has further revealed novel LC3B proximal proteins such as GAPD1 and eIF2D.

## Results

### APEX2-GFP-LC3B labels autophagosomes in HeLa cells

To determine the proteins adjacent to LC3B, we used an APEX2 (Ascorbate Peroxidase 2) proximity labelling approach, originally developed by Rhee et al. (50). We generated HeLa cells stably expressing the APEX2-GFP-LC3B or GFP-LC3B construct (**Fig. 1A**) as shown by western blot (**Fig. 1B**). The localisation of APEX2-GFP-LC3B was investigated by confocal microscopy. Several LC3B puncta were easily detected in cells without any treatment (**Fig. 1C top panel**). Their number was greatly increased upon a three hour treatment with the V-ATPase inhibitor Bafilomycin A1, which impairs lysosomal acidification, leading to an accumulation of autophagosomes (51). LC3B puncta were slightly increased upon treatment with Torin 1, which inhibits mTOR activity, confirming that the APEX2-GFP-LC3B construct allows an increased formation of autophagosomes and an intact autophagic flux (52). Treatment with both drugs led to an increase of LC3B-positive puncta and clear colocalisation with the late endosome/lysosomal marker LAMP1, indicating an autophagosome accumulation (**Fig. 1C**).

**Fig. 1:**
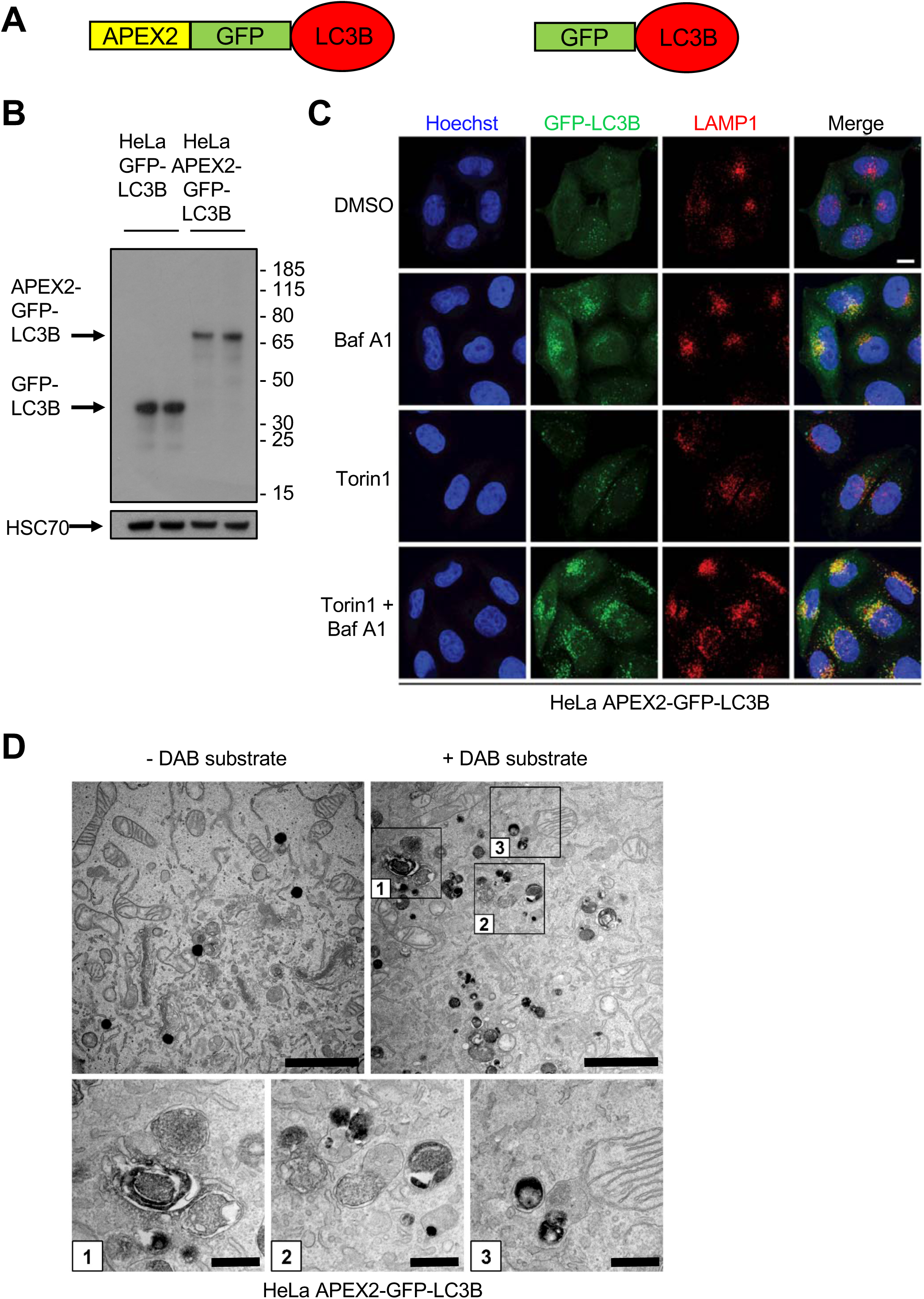
APEX2-GFP-LC3B labels autophagosomes. **(A)** Schematic of APEX2-GFP-LC3B and GFP-LC3B constructs. **(B)** Expression of LC3B and the loading control HSC70 was evaluated in HeLa GFP-LC3B and HeLa APEX2-GFP-LC3B lysates by western blot. **(C)** Confocal sections of HeLa APEX2-GFP-LC3B cells treated with bafilomycin A1 (baf A1; 10 nM), torin 1 (250 nM) or a combination of both for 3 hours. GFP-LC3B (green), LAMP1 (red) and Hoechst (blue). Colocalisation appears as yellow puncta. Scale bar, 10 μm. **(D)** HeLa APEX2-GFP-LC3B cells were treated for 3 hours with 10 nM bafilomycin A1 and 250 nM torin 1 prior to sample processing for electron microscopy with or without DAB substrate. Scale bar, 2 μm. Higher magnifications of boxes 1, 2 and 3 are shown below. Scale bar, 0.5 μm.

In the presence of H_2_O_2_, APEX2 peroxidase tags triggers the polymerization of the peroxidase substrate, 3,3′-diaminobenzidine (DAB), into a localised precipitate that gives transmission electron microscopy (TEM) contrast after treatment with OsO_4_. This allows imaging of cellular compartments, with a very good preservation of ultrastructure and a tighter localisation of the TEM stain than with horseradish peroxidase (HRP) (53). Such staining was observed in HeLa APEX2-GFP-LC3B cells following these treatments. Higher magnification electron micrographs revealed that structures positive for APEX2-GFP-LC3B share features of autophagosomes, including double membranes and engulfed cytoplasmic content (**Fig. 1D**).

These results confirm that APEX2-GFP-LC3B specifically labels autophagosomes. Such an approach would likely be useful to improve the ultrastructural imaging of autophagosomes and could be used for proximity proteome profiling.

### APEX2 workflow and validation of the method

**Fig. 2A** illustrates the workflow of our study aimed at determining the LC3B proximity proteome. HeLa APEX2-GFP-LC3B and HeLa GFP-LC3B cells, cultured for 24 hours in full medium (containing 10% FBS), were treated with biotin phenol for 30 minutes and H_2_O_2_ for 1 minute, allowing APEX2 to generate biotin-phenoxyl radicals that covalently tag proximal endogenous proteins within 20 nm (49) (**Fig. 2A panel 1**). The technique was validated in two ways, both by immunofluorescence microscopy and by western blot (**Fig. 2A panel 2**). Confocal microscopy analysis detected streptavidin-546 punctate labelling in HeLa APEX2-GFP-LC3B cells treated with biotin phenol but only in the presence of H_2_O_2_. Importantly, strepatvidin-546 puncta colocalised with or were in proximity to LC3B puncta (**Fig. 2B**). Biotinylated proteins were pulled-down with neutravidin and detected by western blot using streptavidin-HRP. Treatment with H_2_O_2_ increased the amount of biotinylated protein in HeLa APEX2-GFP-LC3B cells but not in HeLa GFP-LC3B cells (**Fig. 2C**). **Panels 3 and 4 in Fig. 2A** summarise the approach for the larger scale proximity proteomics experiment and hit validation, described in the next figures.

**Fig. 2:**
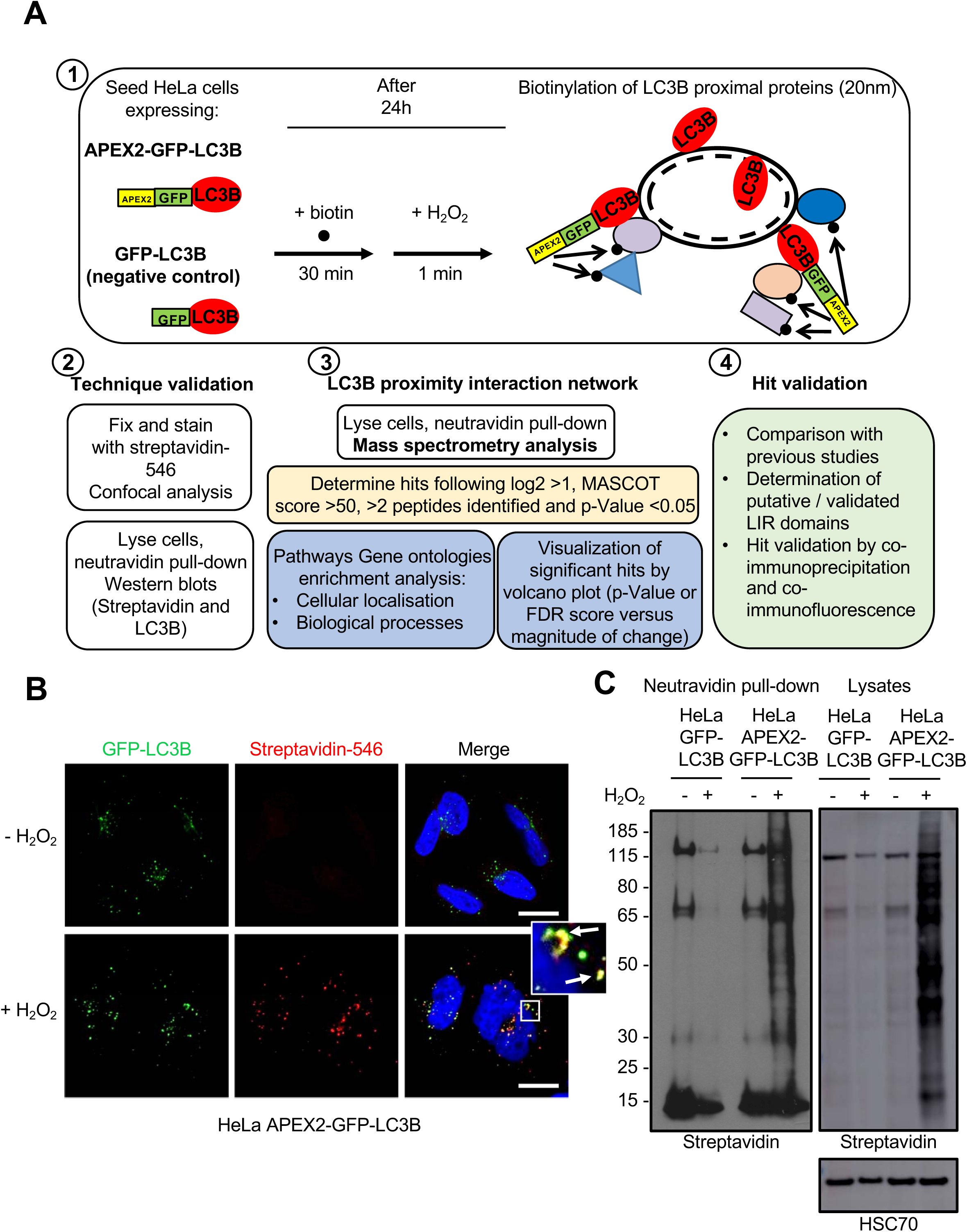
APEX2 workflow and validation of the method. (**A**) Diagram showing the experimental workflow. 1) HeLa cells expressing APEX2-GFP-LC3B or GFP-LC3B were incubated with biotin phenol for 30 minutes and with H_2_O_2_ for 1 minute, allowing biotinylation of proteins in proximity to APEX2-GFP-LC3B. 2) The technique was validated by immunofluorescence and western blot with streptavidin isolation. 3) The experiment was scaled up for proteomic analysis. Here biotinylated proteins were pulled-down with neutravidin and analysed by mass spectrometry followed by a range of bioinformatic and statistical analyses. 4) Hits were validated based on literature research, interrogation of the LIR database and further technical validation for selected proteins. (**B**) Confocal sections of HeLa APEX2-GFP-LC3B cells incubated with biotin phenol for 30 minutes with or without a subsequent 1 minute incubation with H_2_O_2_. GFP-LC3B (green), Streptavidin-546 (red) and DAPI (blue). Colocalisation appears as yellow puncta. Scale bar, 10 μm. (**C**) Levels of streptavidin-HRP evaluated by western blot in HeLa GFP-LC3B and HeLa APEX2-GFP-LC3B neutravidin pull-down eluates following the APEX2 assay (left) or corresponding lysates (right). Cells were incubated with biotin phenol for 30 minutes and with or without H_2_O_2_ for 1 minute.

### APEX2 experiment: 407 positive hits were detected in proximity to LC3B

We scaled up the experiment in order to detect the proteins proximal to LC3B. Both cell lines were incubated with biotin phenol for 30 minutes and H_2_O_2_ for 1 minute. HeLa GFP-LC3B cells were used as a negative control for the APEX2 reaction. Following the reaction, cells were lysed and biotinylated proteins were isolated with a neutravidin pull-down. Before mass spectrometry analysis, the enrichment of biotinylated proteins in the lysates subjected to the neutravidin pull-down were verified by western blot using the streptavidin-HRP and an LC3B antibody. A much more intense smear of biotinylated proteins (**Fig. 3A**) and LC3B (**Fig. 3B**) was detected in HeLa APEX2-GFP-LC3B compared to HeLa GFP-LC3B.

**Fig. 3:**
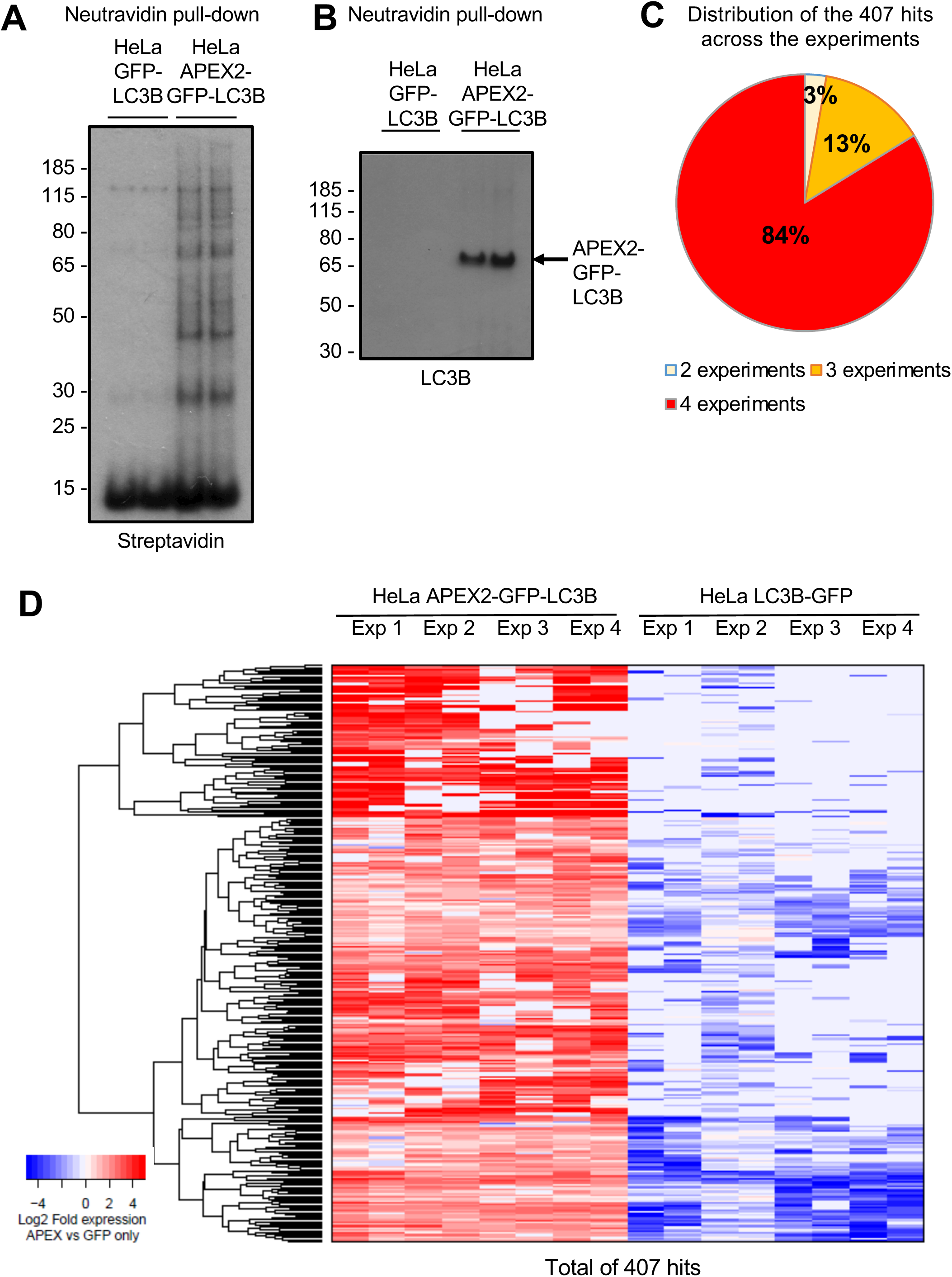
407 hits in proximity to APEX2-GFP-LC3B were detected by proteomics. (**A, B**) Before the mass spectrometry analyses, levels of biotinylated proteins detected by streptavidin-HRP (**A**) and LC3B (**B**) were evaluated by western blot in HeLa GFP-LC3B and HeLa APEX2-GFP-LC3B neutravidin pull-down eluates following the APEX2 assay. Cells were incubated with biotin phenol 30 minutes and H_2_O_2_ for 1 minute. The experiment was performed in duplicates. (**C**) Pie graph showing the percentage of the 407 proteomic hits overlapping between 2, 3 or 4 experiments. (**D**) Heat map representation of the 407 hits in HeLa APEX2-GFP-LC3B cells and HeLa GFP-LC3B cells in each of the four experiments. Proteins were selected if log2 HeLa APEX2-GFP-LC3B compared to HeLa GFP-LC3B >1, MASCOT score >50, >2 peptides identified and p-Value <0.05.

The samples were processed for an on-bead digest and subsequently analysed by liquid chromatography tandem mass spectrometry (LC-MS/MS). Four experiments were performed (see details in Methods). Proteins were considered to be hits when they met the following criteria in at least two independent experiments, comparing results obtained with HeLa APEX2-GFP-LC3B versus HeLa GFP-LC3B: log2fold >1, MASCOT score >50, >2 peptides identified and unadjusted p-Value <0.05 (see Methods). Such analyses detected 407 proteins in proximity (distance less than 20 nm) to APEX2-GFP-LC3B, with 237 of these having an FDR <0.25. Eighty-four percent of the hits were found in the four experiments and 97% were found in at least three experiments, demonstrating the reproducibility and robustness of our results (**Fig. 3CD**).

### The 407 LC3B proximity proteins cover a broad range of pathways and cell localisations

We ran a gene ontology (GO) analysis to identify the subcellular localisation of the 407 hits. We selected the GO terms with a p-Value <0.05 and grouped the hits into seven categories: cytoplasm/cytosol, cytoskeleton/cell junction, centrosome/spindle, endosome/trafficking, nucleus, endoplasmic reticulum (ER)/Golgi apparatus and pre-autophagosome/autophagosome (**Fig. 4A and Figs. S1 and S2**).

**Fig. 4:**
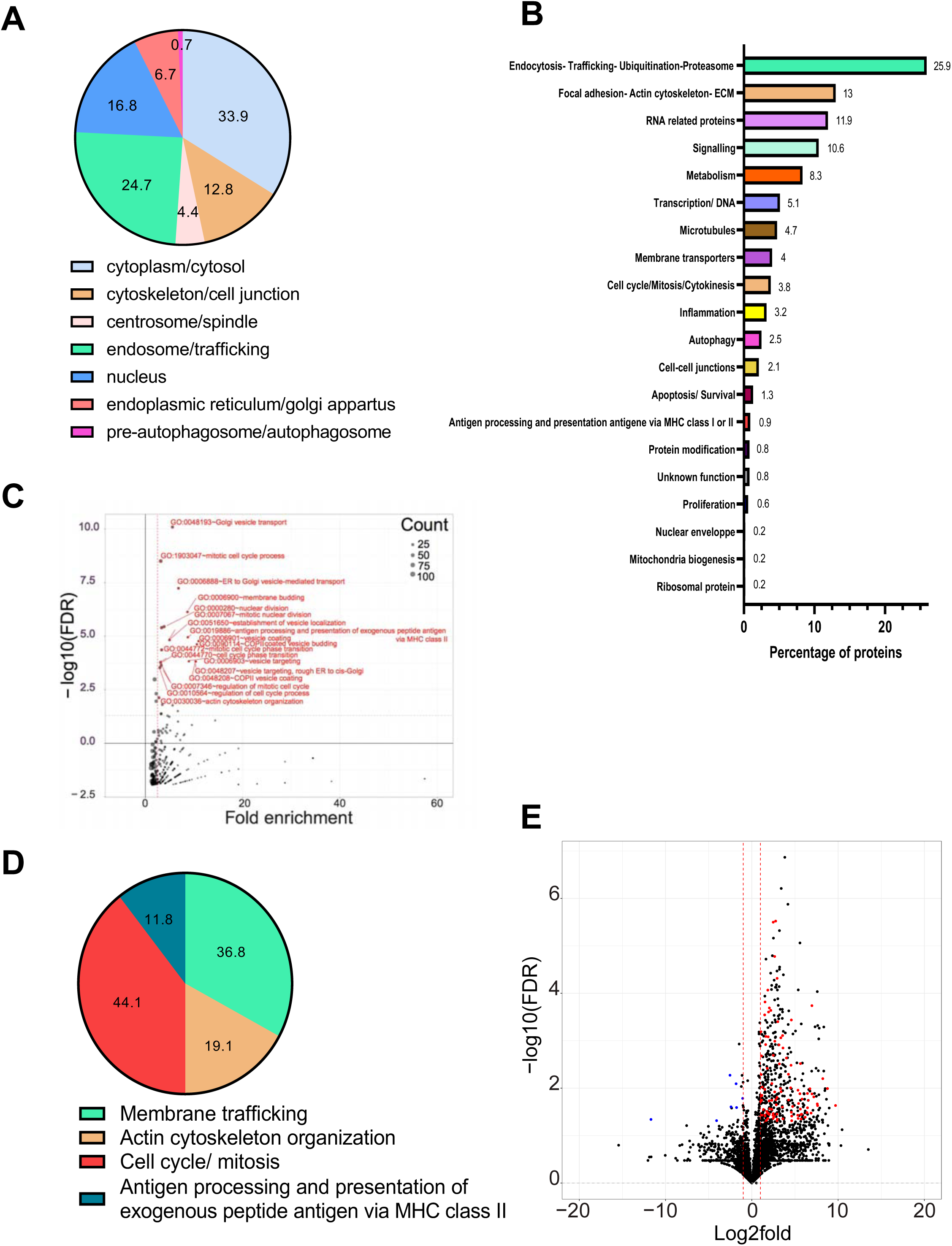
The 407 hits grouped in localisations, pathways, gene ontology enrichment and high confidence hits. (**A)** Pie graph representation of different localisations found by gene ontology (GO) analysis. The percentage of proteins in each localisation is indicated. (**B)** Bar graph showing the different pathways found by an analysis with DAVID software (https://david.ncifcrf.gov/) completed with a search by PubMed and UniProt. The percentage of proteins in each pathway is indicated. (**C, D**) Enrichment analysis of “biological processes” GO analysis. (**C**) Volcano Plot analysis of “biological processes” GO analysis. Proteins were selected following p-Value <0.05 and MASCOT score >50. The names and the red points are for gene ontologies with FDR <0.1 and fold enrichment >2.5. (**D**) Pie graph representation of the different “biological processes” genes found by GO and grouped into four categories. The percentage of proteins in each group is indicated. (**E**) Volcano plot revealing the “high-confidence hits” representing the FDR in function of the Log2fold of the hits obtained in HeLa APEX2-GFP-LC3B cells compared to HeLa GFP-LC3B cells. Hits with a Mascot score >50, >2 peptides and a FDR score <0.05 were selected.

Using DAVID Bioinformatics Resources 6.8 (https://david.ncifcrf.gov/ (54, 55)) to interrogate the KEGG pathways, followed by manual annotations using PubMed (https://pubmed.ncbi.nlm.nih.gov/) and UniProt (https://www.uniprot.org/ (56)), we determined that the 407 identified proteins are involved in a broad range of cellular pathways. These include “Endocytosis, trafficking, ubiquitination or proteasome”, “RNA related proteins”, “Cell cycle, mitosis and cytokinesis” and “Autophagy” (**Figs. 4B and S3**).

To determine which pathways were significantly enriched, we ran a “biological processes” GO analysis of the 407 hits. A subset of 109 proteins were enriched in 18 “biological processes” GO terms, with an FDR <0.1 and a fold enrichment >2.5 (**Figs. 4C and S4)**. We organized these 18 GOs into the four following categories: membrane trafficking (36.8% of the hits), actin cytoskeleton organization (19.1%), antigen processing and presentation of exogenous peptide antigen via MHC class II (11.8%) and cell cycle/mitosis (44.1%) (**Figs. 4D and S5**).

To identify high confidence proximal proteins, we presented all 407 hits in a volcano plot, showing statistical significance (FDR score or p-Value) versus magnitude of change (fold change) between proteins detected in proximity to APEX2-GFP-LC3B and GFP-LC3B. Proximity proteins represented in these plots have a Mascot score >50, >2 peptides and a FDR score <0.05 (**Fig. 4E**) or p-Value <0.05 (**Fig. S6B**). Eighty-eight proximity proteins were detected (**Fig. S6A**).

### Thirty-four percent of the LC3B proximity proteins are already related to autophagy

To validate our approach, we investigated which of the 407 LC3B proximity proteins have already been reported to play a role in autophagy or to be associated to the autophagosome and/or LC3.

This was determined through bioinformatic DAVID Pathway and GO localisation analyses (14 hits, **Figs. 5A in yellow circle and S8**) and through literature research of the high confidence hits revealed by the volcano plot representation (35 hits, **Figs. 5A in green circle and S8**). Further hits were found to overlap with four previous proteomic studies aiming at detecting proteins present in autophagosomes (Dengjel J. et al. (57), Mancias J. D. et al. (58), Gao W. et al. (59), Le Guerroué F. et al. (60)). In these studies, cells were treated in various ways to induce accumulation of autophagosomes. Quantitative proteomics was performed from purified autophagosomes (fractionation or immuno-purification) (57–59) or following an APEX2-GFP-LC3B pull-down from inside autophagosomes only (60). One hundred forty-eight of our 407 hits overlapped with at least one of these proteomic studies, including 68 further reported to play a role in autophagy or in its regulation (**Figs. 5 in grey circle, S7 and S8**). A literature search determined 41 additional hits related to autophagy that were not found with the above-cited methods (**Figs. 5A inside red circle and S8**).

**Fig. 5:**
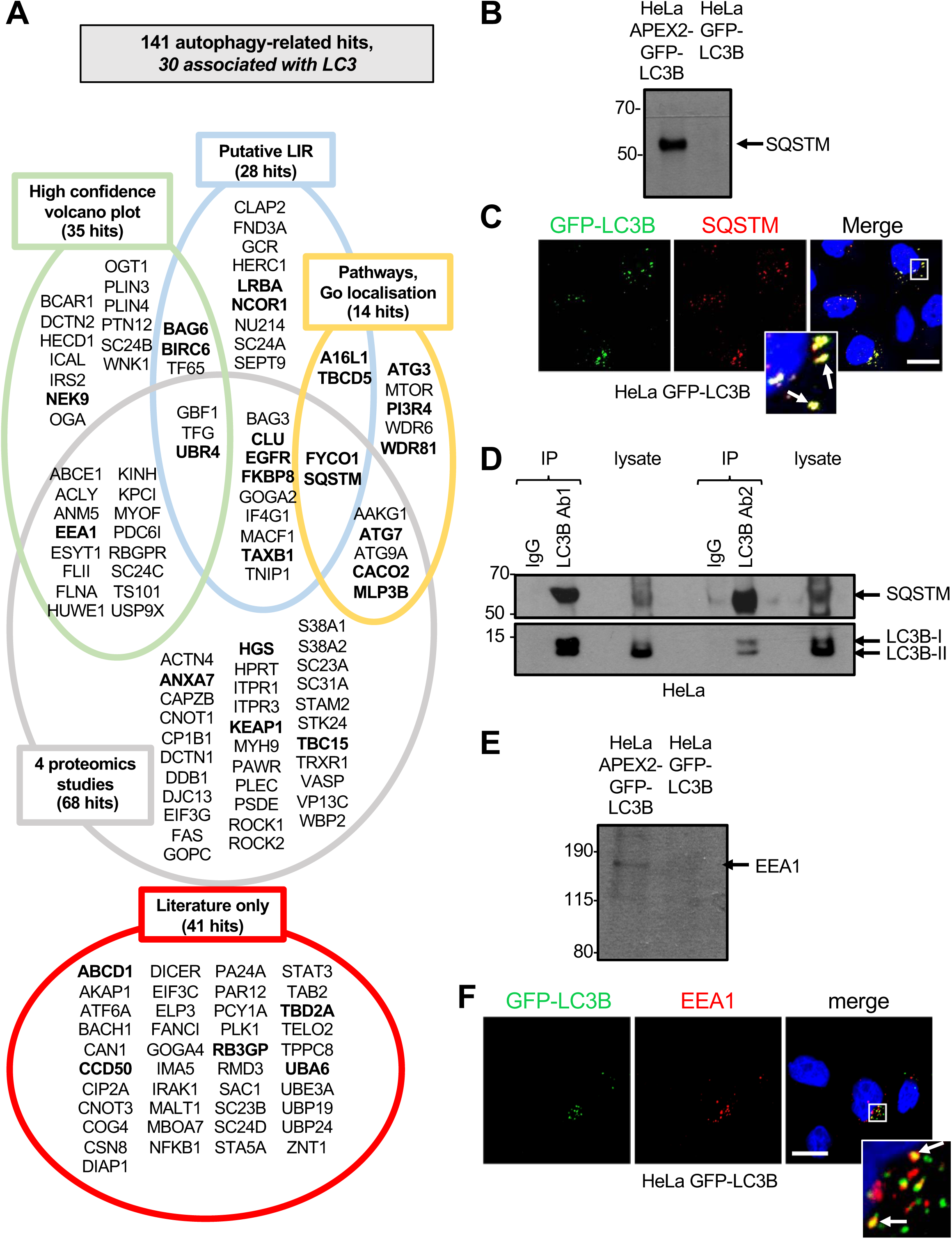
Autophagy-related hits. (**A**) List of the 141 autophagy-related hits. Proteins in bold have been shown associated with LC3 (through co-immunoprecipitation, pull-down or co-immunofluorescence). Green circle: high confidence hits revealed by the volcano plot representation in **Fig. 4E**. Blue circle: containing a putative LIR. Yellow circle: present in pathways of autophagy (DAVID search) or in the GO localisation in autophagosome. Grey circle: present in at least one of the four proteomics studies analysed (57–60). Red circle: present in literature only. (**B, C, D**) Validation of LC3B-SQSTM proximity. (**B**) Western blot for SQSTM from neutravidin pull-down eluates following the APEX2 assay in HeLa APEX2-GFP-LC3B and HeLa GFP-LC3B cells. (**C**) Confocal sections of HeLa GFP-LC3B cells. GFP-LC3B (green), SQSTM (red) and DAPI (blue). Colocalisation appears in yellow. Scale bar, 10 μm. (**D**) Western blots for LC3B and SQSTM following immunoprecipitation of endogenous LC3B in HeLa cells performed in duplicate with an Abcam antibody (Ab1, left side of the blot) and a Cell Signalling Technology antibody, clone D11 (Ab2, right side of the blot) or corresponding IgG control. Total levels of LC3B-I and II and SQSTM in the cell lysates are also shown. (**E, F**) Validation of LC3B-EEA1 proximity. (**E**) Western blot for EEA1 from neutravidin pull-down eluates following the APEX2 assay in HeLa APEX2-GFP-LC3B and HeLa GFP-LC3B cells. (**F**) Confocal sections of HeLa GFP-LC3B cells. GFP-LC3B (green), EEA1 (red) and DAPI (blue). Colocalisation appears in yellow. Scale bar, 10 μm.

We further interrogated the iLIR database (https://ilir.warwick.ac.uk (18)), containing all the putative canonical LIR-containing proteins identified *in silico* in the proteomes of eight model organisms, using the iLIR server (http://repeat.biol.ucy.ac.cy/iLIR/ (61)). Sixty-nine of our 407 hits were found in the iLIR database. Twenty-eight of these have a reported role in autophagy (**Figs. 5A inside blue circle, S8 and S9**), including 13 hits shown to associate (through co-immunoprecipitation or GST pull-down experiments) or colocalise with LC3B: ATG16L1 (A16L1), BAG6, BIRC6, CLU, EGFR, FKBP8, FYCO1, LRBA, NCOR1, SQSTM, TAX1BP1 (TAXB1), TBC1D5 (TBCD5), UBR4 (**Figs. 5A, inside blue circle in bold, S8 and S9**).

Within our autophagy hits, several core autophagy proteins or well-established regulators such as ATG3 (62), ATG7 (63), ATG9A (8), ATG16L1 (A16L1) (16), CALCOCO2 (CACO2) (64), FYCO1 (21), MTOR (65) and SQSTM (66), as well as LC3B (MLP3B) (10), were found (**Fig. 5A**). SQSTM/p62, known as the major receptor for selective autophagy (67), interacts directly with LC3B through its LIR (21), even in basal cell conditions (68). The presence of SQSTM was detected in the neutravidin pull-down (**shown in Fig. 3**) from HeLa APEX2-GFP-LC3B cells but not HeLa GFP-LC3B cells (**Fig. 5B**). SQSTM colocalised with GFP-LC3B puncta (**Fig. 5C**) in HeLa GFP-LC3B cells, ruling out non-specific detection of SQSTM with the APEX2 tag. Furthermore, SQSTM co-immunoprecipitated with LC3B in HeLa cells (expressing endogenous LC3B), using two distinct antibodies previously validated to correctly detect autophagosomes (69, 70) (**Fig. 5D**).

One of our high confidence hits was EEA1 (Early Endosome Antigen 1). This was revealed by the volcano plot representation and also detected in one of the four proteomic studies (57) (**Fig. 5A in intersection between green and grey circles, Fig. S8**). It is recruited to endosomal membranes by binding the phospholipid phosphatidylinositol 3-phosphate (PI(3)P) through its C-terminal FYVE domain (71) where it binds to its effector Rab5 (72, 73). It promotes sorting from the early endosome and early-late endosome fusion (74). Autophagic vacuoles can also fuse with EEA1-positive early endosomes, generating amphisomes (75). Moreover EEA1-positive endosome and autophagic vacuole fusion have been shown to be required for autophagosome maturation and autophagy (76). We confirmed the presence of EEA1 in the neutravidin pull-down (**shown in Fig. 3**) from HeLa APEX2-GFP-LC3B cells and not HeLa GFP-LC3B cells (**Fig. 5E**). GFP-LC3B and EEA1 colocalisation was also detected (**Fig. 5F**).

Therefore, at least 141 LC3B proximity proteins, 34.6 % of our total 407 hits, were found to be related to autophagy (**Fig. 5A and Figs. S8 and S10**), including several proteins with putative LIR domains, validating our method. Interestingly 30 of these 141 autophagy-related hits were shown to associate (through co-immunoprecipitation or GST-pull-down experiments) or colocalise with LC3B (**Figs. 5A, in bold and S8**). Our study also revealed that 111 of the 141 autophagy-related hits have no published interaction or colocalisation with LC3 to our knowledge (others than in the four proteomic studies) (**Fig. S8**).

### The endosomal protein GAPD1 has a putative LIR domain and is in proximity to LC3B

Among our 407 hits, 41 have not been reported to associate with LC3 or to be linked to autophagy, but have a putative LIR domain (**Fig. 6A**). Twenty-nine of these proteins were found to be part of “endocytosis or membrane trafficking” in the DAVID Pathway analysis or localised on endosomes in the GO localisation analyses (**Figs. 6B and S11**). “Endosomal trafficking” is one of the four pathways enriched in the GO “biological processes” analysis (**Fig. 4D**). One of these proteins is GAPD1 or GAPVD1 (GTPase-activating protein and VPS9 domain-containing protein 1), also called RAP6 (Rab5-activating protein 6) and GAPex5. As seen in the diagram **Fig. 6C**, obtained by the software STRING, GAPD1 is an endosomal protein that biologically interacts with a number of others such as Rab5A and B. It plays a role in constitutive and regulated endocytosis and is involved in the regulation of receptor-mediated endocytosis (77). GAPD1 was detected by western blot in the neutravidin pull-down from the HeLa APEX2-GFP-LC3B cells and not HeLa GFP-LC3B cells (**Fig. 6D**). We then confirmed GAPD1 colocalisation with GFP-LC3B puncta (**Fig. 6E**). Furthermore, GAPD1 co-immunoprecipitated with endogenous LC3B in HeLa cells (**Fig. 6F**).

**Fig. 6:**
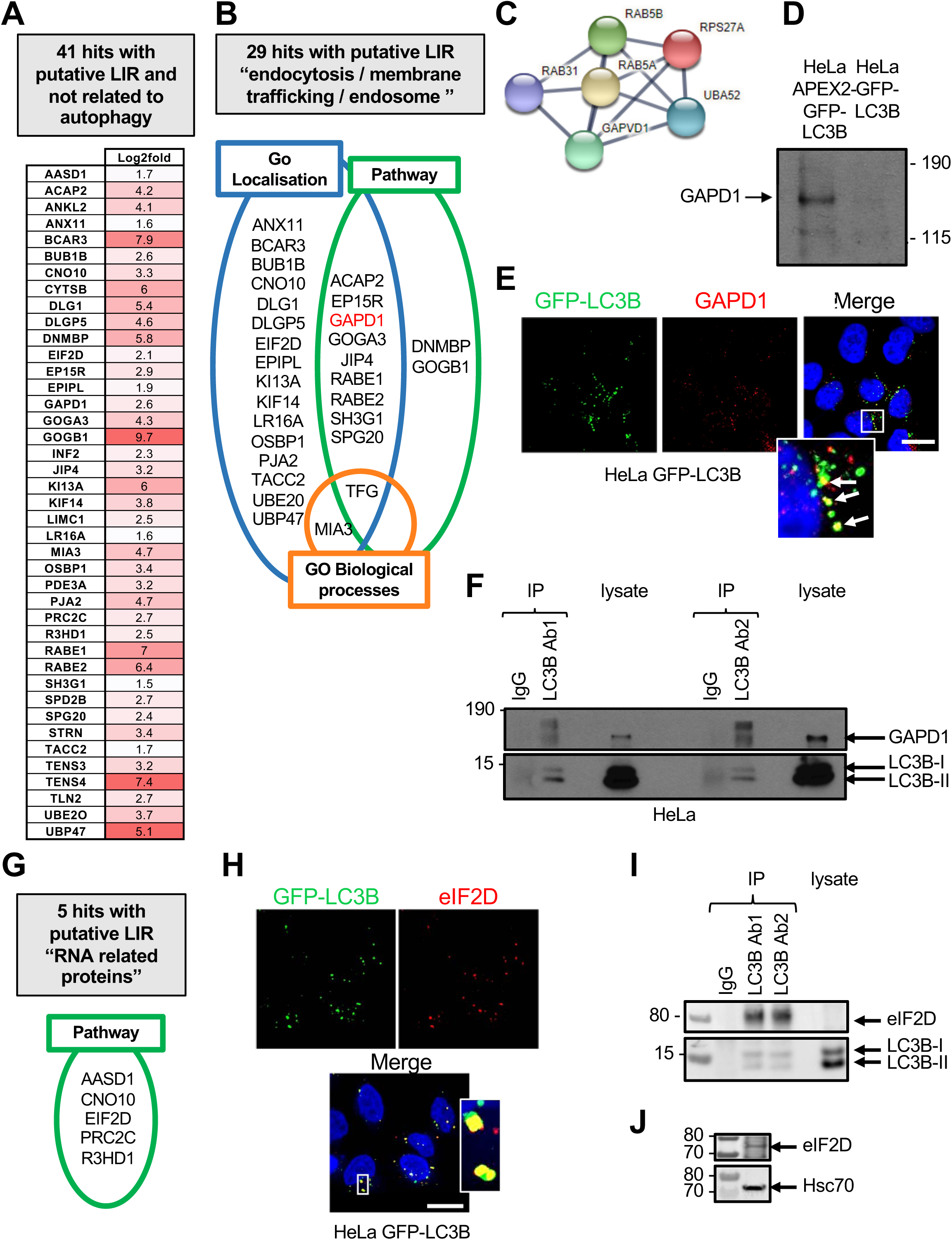
Hits not reported related to autophagy but with putative LIR domains and validation: trafficking pathway and RNA binding/Initiation of translation factors. (**A)** Heat map of the 41 hits not previously shown to be related to autophagy and found in the iLIR database. For each protein, the log2fold HeLa APEX2-GFP-LC3B compared to HeLa GFP-LC3B is indicated with a matching colour gradient. (**B**) List of the hits from (**A**) reported to be involved in endocytosis/membrane trafficking pathways in the DAVID pathway database (green circle), in the “biological processes” GO analysis (orange circle) or localised to endosomes as determined by the GO localisation analysis (blue circle). (**C**) GAPD1 network obtained by STRING software. (**D, E, F**) Validation of LC3B-GAPD1 proximity (**D**) Western blot for GAPD1 from neutravidin pull-down eluates following the APEX2 assay in HeLa APEX2-GFP-LC3B and HeLa GFP-LC3B cells. (**E**) Confocal sections of HeLa GFP-LC3B cells. GFP-LC3B (green), GAPD1 (red) and DAPI (blue). Colocalisation appears in yellow. Scale bar, 10 μm. (**F**) Western blots for LC3B and GAPD1 following immunoprecipitation of endogenous LC3B in HeLa cells performed in duplicate with an Abcam antibody (Ab1, left side of the blot) and a Cell Signalling Technology antibody, clone D11 (Ab2, right side of the blot) or corresponding IgG control. Total levels of LC3B-I and II and GAPD1 in the cell lysates are also shown. (**G**) List of the proteins not previously shown to be related to autophagy and found in the iLIR database (from **Fig. 6A**) and reported to be involved in RNA binding, degradation or translation in the DAVID pathway database. (**H**) Confocal sections of HeLa GFP-LC3B cells. GFP-LC3B (green), eIF2D (red) and DAPI (blue). Colocalisation appears in yellow. Scale bar, 10 μm. (**I**) Western blots for LC3B and eIF2D following immunoprecipitation of endogenous LC3B in HeLa cells performed in duplicate with an Abcam antibody (Ab1, left side of the blot) and a Cell Signalling Technology antibody, clone D11 (Ab2, right side of the blot) or corresponding IgG control. Total levels of LC3B-I and II and eIF2D in the cell lysates are also shown. (**j**) Western blots for eIF2D and HSC70 from a large amount of protein lysate.

### The translation initiation factor eIF2D has a putative LIR domain and is associated with LC3B

Five of the LC3B proximity proteins with putative LIR domains, which have not been previously reported to bind LC3, are part of the category “RNA related proteins” in the DAVID Pathway analysis. These are AASD1, CNO10, EIF2D, PRC2C and R3HD1 (**Figs. 6AG, S3 and S11**). EIF2D is an initiation factor of translation, part of the 43S preinitiation complex. Interestingly, the following hits were not found in the ILIR database, but are also initiation factors: eIF3A, eIF3B, eIF3C, eIF3G, eIF3H, eIF3I (78) (**Fig. S11**). EIF2D colocalised with GFP-LC3B puncta (**Fig. 6H**) and co-immunoprecipitated with endogenous LC3B (**Fig. 6IJ**), validating the APEX2 result. While eIF2A phosphorylation has previously been shown to be necessary for the stimulation of autophagy, it is through transcription of several autophagy genes (SQSTM, LC3B, BECN1, ATG3, ATG7) (79, 80). Therefore, our results, revealing a novel LC3B-eIF2D interaction, suggest a novel relationship between the initiation of translation and LC3B.

## Discussion

In this study, we used peroxidase-mediated proteomic mapping in live cells combined with unlabelled mass spectrometry to determine the proteins in proximity (≤ 20 nm) to LC3B in HeLa cells cultured in full medium. Four hundred and seven LC3B proximity proteins were detected. One hundred forty one of these proteins, or 34.6% of the total hits, have been previously reported as related to the autophagy process or its regulation (**Fig. 5A**), validating our approach. These include several core autophagy drivers or regulators. Sixty-eight of these 141 autophagy-related proteins have been determined to be present in the autophagosome or associated with LC3 in at least one of four previous proteomic studies (57–60). Moreover, 30 of the 141 proteins have already been shown to associate with LC3 through co-immunoprecipitation, GST pull-down or colocalisation studies (**Fig. 5A**). Using co-immunofluorescence analysed by confocal microscopy and co-immunoprecipitation experiments, we validated two of these proteins in our culture conditions: the autophagy receptor SQSTM (or p62) (67) (**Fig. 5BCD**), containing an LIR (18, 21), and the sorting endosome marker EEA1 (72, 75) (**Fig. 5EF**). Twenty-eight of the 141 autophagy-related hits were found in the iLIR database, including 13 with demonstrated LC3B association (**Fig. 5A in bold in the blue circle**), further validating our approach.

Interestingly, for 53 hits found in at least one of the four proteomic studies and shown to play a role in autophagy, there is no association with LC3 further reported in the literature (through co-immunoprecipitation or GST pull-down or colocalisation) (**Fig. S7**). It is, for example, the case for ROCK1 and ROCK2, two key protein kinase regulators of the actin cytoskeleton and cell polarity (81). Furthermore, 69 of the total 407 hits were found in the iLIR database, including 56 proteins that have not, to our knowledge, been shown to be associated with LC3 (through co-immunoprecipitation or GST-pull-down or colocalisation) (**Figs. 6A and S9**). On the other hand, 41 proteins with a putative LIR domain have not been reported to be related to autophagy (**Fig. 6A**). These findings suggest that a direct interaction between some of these proteins and LC3B may regulate an unknown function.

Previous proteomic studies aimed to determine proteins present in the autophagosome and thus were performed from enriched autophagosome from cells that were serum or amino acid-deprived or treated with inhibitors to trigger stress-induced autophagy or inhibit autophagic flux (60, 82). Interestingly, a LC3-APEX2 screen (60) has recently been published, and 54 of our hits are common with this study (**Fig. S7**). This study determined LC3B proximity proteins inside the autophagosome. Our study has a distinct focus from these previous studies, which is to uncover the LC3B global interactome, irrespective of LC3B localisation (inner or outer membrane of the autophagosome or other cell location). We have determined the LC3B interactors from whole cell lysates and in basal cell condition (using cells maintained in full medium and without inhibitors). Thus our experimental conditions allow the detection of proteins in proximity to LC3B potentially involved in the three primary types of autophagy: microautophagy (83), macroautophagy (6), and CMA (84), but also in non-canonical autophagy or even functions unrelated to autophagy (26, 35, 38, 85, 86). Additional LC3-ylated proteins (that we recently described (47)) may also have been uncovered.

Accordingly, we found some novel LC3B interactors, involved in a broad range of pathways and cell localisation (**Fig. 4**). The pathway functions include membrane trafficking, cytoskeleton rearrangement, antigen processing and presentation of exogenous peptide antigens, RNA-related proteins, cell cycle/mitosis/cytokinesis and nuclear proteins. Interestingly, a pool of LC3 has recently been shown to localise to the nucleus and initiate autophagy upon redistribution to the cytoplasm (63, 87–89). Over 16% of the 407 proteins detected in proximity to LC3B have a reported nuclear localisation (**Fig. 4A**), and some could be novel partners or regulators of LC3B nucleo-cytoplasm shuttling. Moreover, LC3 has been reported to associate with nuclear complexes (45, 90).

A substantial proportion of our hits were part of pathways we regrouped under “actin cytoskeleton organisation/focal adhesion/ECM/cell junction” (**Fig. 4**). Among these, two proteins of the alpha-actinin family (ACTN1 and ACTN4) are cytoskeletal actin-binding proteins with an important role in the structure and regulation of cytoskeleton organization and muscle contraction (91). These two proteins were not found in the iLIR database. ACTN4, but not ACTN1, has been described in autophagy (92) although with no interaction with LC3B.

Another group of our identified LC3B interactors were previously reported to be part of the membrane trafficking localisation (24.7%) or pathway (25.9% of the hits) (**Fig. 4AB**). It is one of the four enriched pathway categories in the GO “biological functions” analysis and accounts for 36.8% of the hits enriched (**Fig. 4D**). This could reflect the highly dynamic membrane trafficking which depends on various endomembranes connecting with each other, through maturation or fusion. Nevertheless, 53 of these proteins have a putative LIR domain, including 11 reported associated with LC3, reinforcing the possibility that they are true LC3B interactors (**Fig. S12**). Moreover, 29 proteins with a putative LIR domain and not previously reported to be related to autophagy are membrane trafficking proteins or have been found localised to endomembranes (**Fig. 6B**). We have validated one of these proteins, GAPD1, a regulator of endocytosis that exhibits GEF activity specific for RAB5 and GAP activity specific for RAS (77) (**Fig. 6DEF**).

Eleven-point nine percent of the 407 hits are part of “RNA binding proteins” as detected by the DAVID database (**Fig. 4B**). Several of these proteins have not been reported to be related to autophagy and have a putative LIR domain (**Fig. 6G**). Of these, we have validated the translation factor EIF2D (**Fig. 6HI**). The fact that several other initiation factors of translation, all part of the 43S preinitiation complex (78), were also detected in proximity to LC3B strongly suggest a previously unrecognised relationship between LC3B and this complex.

This study has uncovered the LC3B proximity proteome in cells in normal growth culture. We have discovered a number of novel LC3B proximity proteins, opening the way to a better understanding of LC3B’s role in autophagy and possibly functions not related to autophagy.

## Authors contribution

MN designed, performed and analyzed most of the experiments and wrote the manuscript. AA generated and characterized the constructs and cell lines. FM performed some of the experiments and provided comments on the manuscript. ADD performed the bioinformatic analyses. VR performed and analyzed the proteomics. JB performed and analyzed the electronic microscopy. TN and RK contributed to the experimental design, to the interpretation of the results and provided comments on the manuscript. PC directed the proteomic experiments, performed the related bioinformatic analyses and provided comments on the manuscript. SK conceived and directed the project, designed the experiments and wrote the manuscript.

## Acknowledgements

We thank Sara Farrah Heuss for her assistance in the preliminary APEX2 experiments and Brynna Hoggard for the critical reading of the manuscript.

This work was supported by the Medical Research Council (MR/R009732/1), Rosetrees Trust (M314) and Barts and The London Charity (MGU0511).

## Declaration of interests

The authors declare no competing interest.

## Methods

### Generation of HeLa cells stably expressing APEX2-GFP-LC3B

The APEX2 coding and linker peptide sequence was PCR-amplified from pcDNA3 Connexin43-GFP-APEX2, a gift from Alice Ting (Addgene plasmid # 49385; http://n2t.net/addgene:49385; RRID:Addgene_49385) (93), using the primers 5’-PacI-APEX2 (5’-GACTTAATTAAGCCACCATGGGAAAGTCTTACCCAACTGTG-3’) and 3’-XbaI-linker-APEX2 (5’-GACTCTAGATCCGGAGCCCGAGCCCGAGGTCGAGCCCGAGCCCTTGGCATCAGCAAACCCAAGC-3’). The resulting PCR product was inserted into pLVpuro-CMV-N-EGFP (Addgene plasmid # 122848 ; http://n2t.net/addgene:122848; RRID:Addgene_122848) (94) by restriction-based cloning, between the PacI and XbaI sites, to generate pLVpuro-CMV-N-APEX2-EGFP. Gateway recombination of the destination vector pLVpuro-CMV-N-APEX2-EGFP with the entry clone pDONR223 LC3B WT (Addgene plasmid # 123072; http://n2t.net/addgene:123072; RRID:Addgene_123072) (94) was performed using LR clonase II enzyme mix (ThermoFisher Scientific, 11791020), to generate pLVpuro-CMV-APEX2-EGFP-LC3B. Lentiviral packaging and stable cell line generation was then performed as previously described (94) using pLVpuro-CMV-APEX2-EGFP-LC3B as the transfer plasmid and infecting wild type HeLa cells with the resulting lentiviral particles. A clonal HeLa cell line expressing moderate levels of APEX2-EGFP-LC3B was isolated by limit dilution prior to expansion and use in all experiments described. As a control, wild type HeLa cells were infected in parallel with lentivirus encoding CMV-driven EGFP-LC3B generated previously (94) and expanded as a pool.

### Cell culture

HeLa cells were cultured in Dulbecco’s Modified Eagle’s Medium (DMEM, Gibco) supplemented with 10% fetal bovine serum (FBS, Gibco). For the HeLa cells infected with GFP-LC3B or APEX2-GFP-LC3B lentivirus, puromycin (2μg/ml, Sigma Aldrich) was added in the medium. The cells were grown in a humidified atmosphere with 5% CO_2_ at 37°C.

### Western blot

Cells were lysed in sample buffer (Thermofisher) supplemented with dithiothreietal (DTT, Fisher), sonicated and boiled at 99°C for 5 min. Samples were loaded on 4–12% Novex Bis-Tris gels (Invitrogen). Separated proteins were transferred onto 0.45 μm nitrocellulose transfer membranes (VWR International). Protein loading and transfer quality were checked by staining with Ponceau S. Membranes were blocked in TBS with 4% BSA, then probed with the following primary antibodies: EEA1 (Santa cruz), eIF2D (invitrogen), GAPD1 (Novusbio), HSC70 (Santa Cruz), LC3B (Clone D11, Cell Signaling Technology), and SQSTM (Cell Signaling Technology) at a 1:1000 dilution and Streptavidin HRP (Millipore) at a 1:10,000 dilution. Membranes were incubated overnight at 4°C, followed by a one hour incubation in the appropriate secondary antibodies coupled to peroxidase (1:1000, Biorad). Proteins were detected by enhanced chemiluminescence detection (ECL, GE Healthcare). Densitometric analyses of immunoblots were performed using ImageJ 1.50i (National Institute of Health).

### Immunofluorescence and confocal microscopy

Cells were plated onto coverslips in 24-well plates (50000 cells per well). For most experiments, cells were fixed in 2% paraformaldehyde for 15 minutes. Free aldehydes were quenched with 50 mM NH_4_Cl in PBS for 10 min. Fixed cells were blocked and permeabilized in PBS with 3% BSA and 0.1% Triton X-100 for 15 minutes. Cells were incubated with the following primary antibodies: eIF2D (invitrogen), GAPD1 (RAP6 Novusbio), LC3B (Catalogue number 2775S, Cell Signaling Technology), and SQSTM (Cell Signaling Technology) at a 1:100 dilution 1:100 and streptavidin-546 (Life technologies) at a 1:500 dilution for 30 minutes. Cells were washed three times in PBS, followed by a 20 minutes incubation with the appropriate secondary antibodies coupled to a fluorochrome Alexa 555 (1:500, Life technologies). Cells were washed three times in PBS and once in water and mounted onto slides with Prolong™ Gold Antifade Mountant with DAPI (Molecular probes, Life technologies). Images were acquired using a confocal laser scanning microscope (LSM710; Carl Zeiss, Inc.) equipped with a 63×1,4 NA Plan-Apochromat oil immersion objective. Alexa 488 was excited with the 488 nm line of an Argon laser, Alexa 555 was excited with a 543 nm HeNe laser. For immunofluorescence using the anti-LAMP1 antibody (1:1,000, BD Biosciences, Catalogue number 555798) we used a protocol previously described (94) and imaged with a Leica SPE confocal microscope using a 1.3 NA 63x oil immersion objective.

All image fields were chosen arbitrarily based on DAPI (4,6-diamidino-2-phenylindole) staining and images were taken in unsaturated conditions.

### Electron microscopy

Transmission electron microscopy sample preparation and imaging was performed as previously described (94) with some modifications. The DAB substrate mix was freshly prepared using 3.5 mM DAB (3,3’-Diaminobenzidine tetra–HCl, obtained from TAAB UK) and 0.02% H_2_O_2_ (Sigma) in 50 mM Tris-HCl at a pH of 7.6. Immediately following glutaraldehyde fixation and rinsing in 0.1 M sodium cacodylate, samples were incubated with or without the DAB substrate mix for 3 minutes at room temperature. To stop the reaction, samples were rinsed in 50 mM Tris-HCl buffer followed by 0.1 M sodium cacodylate. Samples were then fixed with osmium and processed as previously described (94), but with the omission of tannic acid and sodium sulphate incubations.

### Co-Immunoprecipitation

Cells were lysed in RIPA buffer (Merck) supplemented with phosphatase and protease inhibitors. Lysates were centrifuged at 10000 rpm for 10 minutes at 4°C. The supernatant was precleared with beads (Protein G sepharose™ 4 flast flow, GE Healthcare) for 1 hour at 4°C on a rotating wheel. Beads were removed and 4 μg LC3B antibody (Abcam or clone D11 Cell Signalling Technology) or IgG was added and rotated overnight at 4°C. The following day, beads were added and rotated for 1 hour at 4°C. Beads were washed 3 times with RIPA and sample buffer supplemented with DTT was added.

### APEX2 experiment

Five million cells were plated in 100 × 17 mm dishes in full media containing 7 μM Heme (Sigma). After 24 hours, the cells were treated with 500 μM biotin tyramide (biotin phenol; Iris Biotech) for 30 minutes at 37°C and exposed to 1 mM freshly prepared hydrogen peroxide for 1 minute. The biotinylation reaction was quenched by 3 washes of a stop solution prepared with 10 mM 6-Hydroxy-2,5,7,8-tetramethylchroman-2-carboxylic acid (Trolox; Sigma), 20 mM sodium Ascorbate and 10 μM sodium azide in PBS. Cells were lysed at 4°C in RIPA buffer supplemented with 10 mM sodium azide and protease inhibitors. Lysates were centrifuged at 14000 rpm for 15 minutes at 4°C and protein concentration determined with a Pierce™ BCA Protein Assay Kit (Life Technologies). High capacity neutravidin beads (Life Technologies) were added to the lysates and rotated overnight at 4°C. Beads were washed 3 times with RIPA. For western blot, sample buffer supplemented with DTT was added. For mass spectrometry, beads were washed with 25 mM ammonium bicarbonate buffer 3 times, centrifuged at 14000 rpm and frozen at −80°C. Three independent experiments were performed, including one experiment in duplicate (called experiment 3 and 4), thus four replicates were produced.

### Mass spectrometry

Proteomics experiments were performed using mass spectrometry as previously described (95, 96). Immunoprecipitated (IP) protein complex beads were digested into peptides using trypsin and peptides were desalted using C18+carbon top tips (Glygen corparation, TT2MC18.96) and eluted using 70% acetonitrile (ACN) with 0.1% formic acid. Dried peptides were dissolved in 0.1% TFA and analysed by ultimate 3000 RSL nanoflow liquid chromatograph coupled on-line to a Q Exactive plus mass spectrometer (Thermo Fisher Scientific). Gradient elution was from 3% to 35% buffer B over 120 minutes at a flow rate of 250 nL/min with buffer A being used to balance the mobile phase. Buffer A was 0.1% formic acid in water and B was 0.1% formic acid in ACN. The mass spectrometer was controlled by Xcalibur software (version 4.0) and operated in the positive ionisation mode. The spray voltage was 1.95 kV and the capillary temperature was set to 255ºC. The Q-Exactive plus was operated in a data-dependent mode with one survey MS scan followed by 15 MS/MS scans. The full scans were acquired in the mass analyser at 375-1500 m/z with a resolution of 70000, and the MS/MS scans were obtained with a resolution of 17500.

MS raw files were converted into Mascot Generic Format using Mascot Distiller (version 2.7.1) and compared to the SwissProt database using the Mascot search daemon (version 2.6.0) with FDR of ∼1% and restricted to human entries. Allowed mass windows were 10 ppm and 25 mmu for parent and fragment mass to charge values, respectively. Variable modifications included in searches were oxidation of methionine, pyro-glu (N-term) and phosphorylation of serine, threonine and tyrosine. The mascot result (DAT) files were extracted into Excel files for further normalisation, quantitative label-free analysis and statistical analysis as described previously (49, 96).

### Bioinformatic analyses

#### Statistical analyses

Statistical analyses were performed within the R (v3.6.1) statistical computing environment (98). Following sum scaling to normalise the protein quantification data, missing values were replaced by the minimum observed value within each sample. For differential analysis, proteins were first filtered on identification confidence criteria (Mascot score >50, >2 detected peptides per protein; in APEX2 only comparisons, any proteins detected in GFP controls were removed) and then the log 2 transformed average fold change was calculated between the conditions. Unpaired, two-tailed t tests were used to assess significance in the proteomics data. Where applicable, p-Values were adjusted for multiple testing using the Benjamini-Hochberg method (99). The analyses were visualized using a combination of individual R packages, namely: ggplot2, gplots, reshape2, Hmisc, readXL and ggrepel (100–105).

#### Gene ontology analysis

For gene ontology analysis, proteins differentially expressed between conditions (at p-Value <0.05) were grouped into gene ontologies as annotated in UniProt and DAVID informatics curated databases (Biological processes/cellular compartment/molecular function) (54–56). To infer ontology enrichment across sets of samples, a hypergeometric test was used followed by Benjamini-Hochberg multiple testing correction.

For literature validation analysis, supplementary data from four studies (57–60) that had previously performed autophagy-associated proteomics studies were extracted. These datasets were merged and matched against the APEX2-enriched proteins identified in this study that met the following criteria: ≥2 Log2 fold change in abundance compared to GFP controls, with a Mascot score >50.

#### STRING protein-protein interaction (PPI) analyses

STRING (Search Tool for the Retrieval of INteracting Genes) V11 (https://string-db.org/) was used to study the PPI network. The settings “multiple proteins” and organism “Homo sapiens” were selected (106).

#### DAVID analyses

DAVID Bioinformatics Resources 6.8, NIAID/NIH (https://david.ncifcrf.gov/) was used to identify the KEGG (Kyoto Encyclopedia of Genes and Genomes) pathways. KEGG is a collection of databases. The settings “UniProt ID” and “gene list” were selected (55).

## Notes

### Competing Interest Statement

The authors have declared no competing interest.

