## Supplementary Figures for "Proximity interactome of LC3B in normal growth conditions"

#### Supplementary Figure Legends

**Fig. S1 (related to Fig. 4A): GO Localisations of the 407 hits**

Bar graphs showing the significantly enriched GO localisations grouped in seven categories: cytoplasm/cytosol, cytoskeleton/cell junction, centrosome/spindle, endosome/trafficking, nucleus, endoplasmic reticulum/golgi apparatus, pre-autophagosome/autophagosome. The -log10 p-Value is represented.

**Fig. S2 (related to Fig. 4A): Hits present in the GO Localisations**

Heat map listing the 407 hits grouped in the seven different GO localisation categories (**Supplementary Fig. 1**). For each protein, the log2fold HeLa APEX2-GFP-LC3B compared to HeLa GFP-LC3B is indicated with a matching colour gradient.

**Fig. S3 (related to Fig. 4B): Distribution of the 407 hits in pathways (DAVID analysis)**

Heat maps listing the 407 hits in the indicated pathways as determined by DAVID software analysis (https://david.ncifcrf.gov/) completed with a PubMed and UniProt search. For each protein, the log2fold HeLa APEX2-GFP-LC3B cells versus HeLa GFP-LC3B is indicated with a matching colour gradient.

**Fig. S4 (related to Fig. 4CD): The four categories of Biological processes**

Bar graphs showing the significantly enriched "biological processes" GO analysis grouped in four categories: membrane trafficking, actin cytoskeleton organization, cell cycle/mitosis and antigen processing and presentation of exogenous peptide antigens via MHC class II. The -log10 p-Value is represented.

**Fig. S5 (related to Fig. 4CD): Distribution of the hits enriched following GO "biological processes" analysis**

Heat map listing, in alphabetical order, the hits from the four enriched categories of GO "biological processes" **(Supplementary Fig. 4).** For each protein, the log2fold HeLa APEX2-

GFP-LC3B cells compared to HeLa GFP-LC3B is indicated with a matching colour gradient. They are grouped in proteins previously shown (left), or not shown (right), to be related to autophagy and whether an association with LC3 (via co-immunoprecipitation or pull-down or co-immunofluorescence) has been reported.

**Fig. S6 (related to Fig. 4E): High confidence hits**

(**A**) Heat map listing, in alphabetical order, the high confidence hits displayed in the volcano plot analysis (presented in **Fig. S6B**). They are grouped as proteins previously shown (left), or not shown (right), to be related to autophagy and whether an association with LC3 (via co-immunoprecipitation or pulldown or co-immunofluorescence) has been reported. For each protein, the log2fold HeLa APEX2-GFP-LC3B compared to HeLa GFP-LC3B is indicated with a matching colour gradient.

(**B**) Volcano plot revealing the "high-confidence hits" representing the p-Value (in Log10) in function of the Log2fold of the hits obtained in HeLa APEX2-GFP-LC3B cells compared to HeLa GFP-LC3B cells. Hits with a Mascot score >50, >2 peptides and a FDR score <0.05 were selected.

**Fig. S7 (related to Fig. 5A): Comparison with four previous studies**

Heat map listing, in alphabetical order, the hits that are common to proteins previously found in the autophagosome or in proximity to LC3 in the four following proteomic studies: Le Guerroué et al., 2017; Dengjel et al., 2012; Gao et al., 2010 and Mancias et al., 2014. They are grouped as proteins previously shown (left), or not shown (right), to be related to autophagy and whether an association with LC3 (via co-immunoprecipitation or pull-down or co-immunofluorescence) has been reported. For each protein, the log2fold HeLa APEX2-GFP-LC3B compared to HeLa GFP-LC3B is indicated with a matching colour gradient.

**Fig. S8 (related to Fig. 5A): Autophagy-related hits**

Heat map listing, by alphabetical order, the 141 hits related to autophagy (as determined by literature research). For each protein, columns, from the left to the right, indicate: the log2fold HeLa APEX2-GFP-LC3B compared to HeLa GFP-LC3B with a matching colour gradient; a reported association with LC3 (via coimmunoprecipitation or pull-down or co-immunofluorescence); presence of a putative LIR; presence in the volcano plot analysis revealing high confidence hits; presence in the four indicated proteomic studies; presence in the autophagy pathway (DAVID analysis)/autophagosome localisation (GO analysis); presence in the “biological processes” GO analysis. The hits written in bold were not found associated to LC3 (through co-immunoprecipitation, pull down or co-immunofluorescence studies) and those further highlighted in orange were present in at least one of the four proteomic studies.

**Fig. S9 (related to Fig. 5A): Hits found in the iLIR database and related to autophagy**

Heat map listing, in alphabetical order, hits which are in the iLIR database (https://ilir.warwick.ac.uk) and related to autophagy. They are grouped as proteins with a reported association with LC3 (via co-immunoprecipitation or pull-down or co-immunofluorescence, left) or not (right). For each protein, the log2fold HeLa APEX2-GFP-LC3B compared to HeLa GFP-LC3B is indicated with a matching colour gradient.

**Fig. S10 (related to Fig. 5A): autophagy-related hits**

Biological network of the 141 hits previously shown to be related to autophagy obtained by STRING software (listed in **Fig. S8**).

**Fig. S11 (related to Fig. 6): 42 hits not previously shown to be related to autophagy or associated with LC3 but with a putative LIR domain and part of membrane trafficking, RNA-related proteins or cell cycle.**

Heat map listing, in alphabetical order, 42 hits previously not shown to be related to autophagy or associated with LC3 but present in the iLIR database and part of trafficking, cell cycle or RNA related proteins as determined by GO “biological process” analysis (orange columns), GO localisation analysis (blue columns) and DAVID pathway (green columns) analyses. For each protein, the log2fold HeLa APEX2-GFP-LC3B compared to HeLa GFP-LC3B is indicated with a matching colour gradient.

**Fig. S12: 53 hits with a putative LIR domain and part of membrane trafficking.**

Heat map listing, in alphabetical order, 53 hits present in the iLIR database and part of trafficking as determined by GO “biological process” analysis, GO localisation analysis and DAVID pathway analyses. For each protein, the log2fold HeLa APEX2-GFP-LC3B compared to HeLa GFP-LC3B is indicated with a matching colour gradient and reported associations with LC3 are indicated.

### Figure S1 (related to Figure 4A)

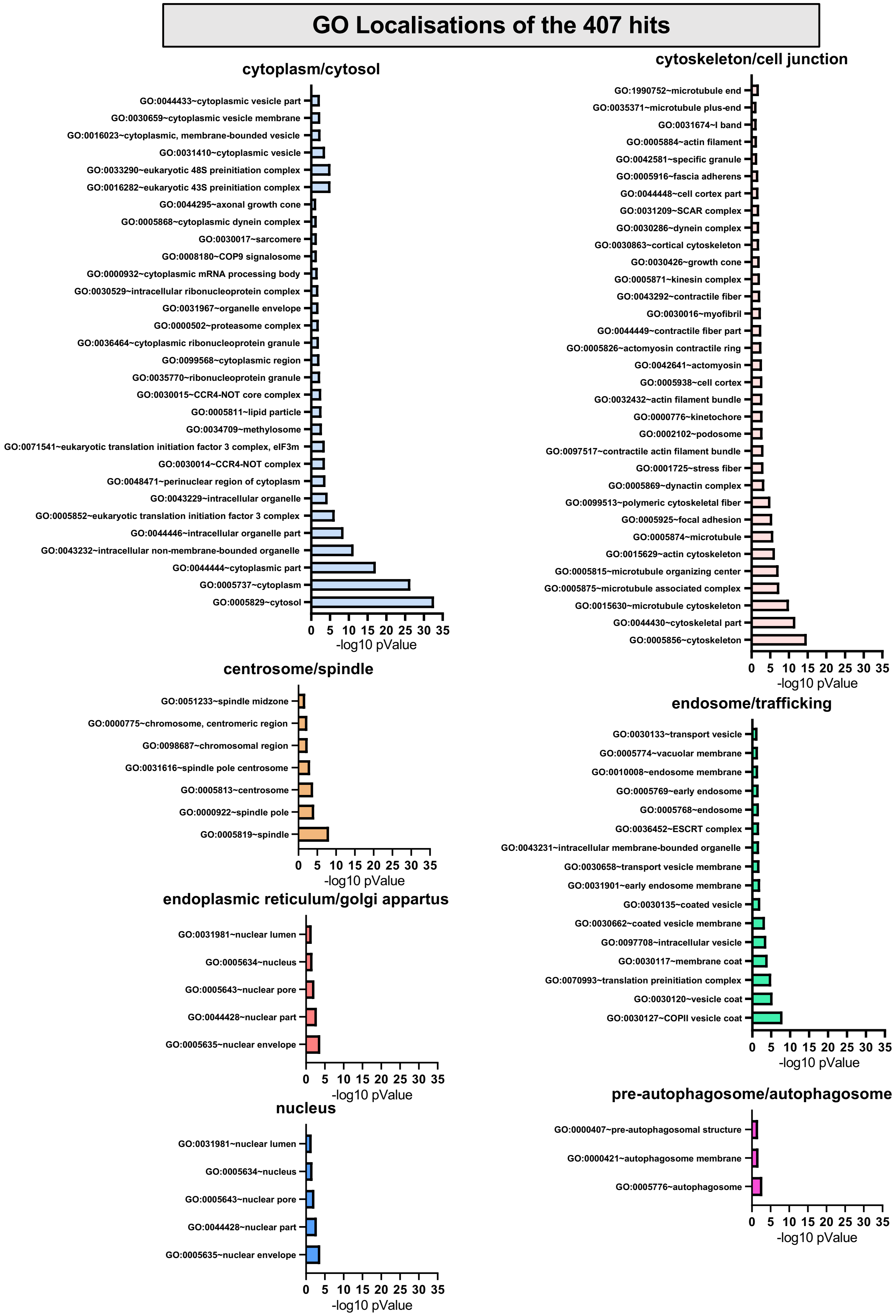

### Table S2 (related to Figure 4A)

#### Hits present in the GO Localisations

| | log2fold | cytoplasm/ cytosol | cyto-skeleton/ cell junction | centro-some/ spindle | endosome/ trafficking | nucleus | endoplasmic reticulum/ Gol apparatus | pre-autophagosome/ autophagosome |
|---|---|---|---|---|---|---|---|---|
| A16A1 | 5.5 | | | X | | | | |
| A16L1 | 3.0 | X | X | | X | | | X |
| AAKG1 | 1.9 | X | | | X | X | | |
| AASD1 | 1.7 | X | | | | | | |
| ABCD1 | 1.2 | X | | | | | | |
| ABCE1 | 1.5 | X | | | X | | | |
| ABI1 | 8.6 | X | X | | X | X | | |
| ABL2 | 4.3 | X | X | | | | | |
| ACAP2 | 4.2 | X | | | X | | | |
| ACLY | 1.8 | X | | | X | X | | |
| ACPH | 4.4 | X | | | X | X | | |
| ACTN1 | 3.4 | X | X | | X | | | |
| ACTN4 | 4.5 | X | X | | X | X | | |
| AHNK | 1.2 | X | X | | X | X | | |
| AIMP1 | 1.3 | X | | | X | X | X | |
| AK17A | 2.1 | X | | | X | X | | |
| AKAP1 | 7.6 | X | | | X | | | |
| ALDOA | 1.2 | X | X | | X | X | | |
| ALG1 | 2.3 | X | | | X | | X | |
| AMPD2 | 2.9 | X | | | | | | |
| ANKL2 | 4.1 | X | | | X | | X | |
| ANKY2 | 3.6 | X | | X | X | | | |
| ANLN | 2.6 | X | X | | X | X | | |
| ANM1 | 1.4 | X | | | X | X | | |
| ANM5 | 2.6 | X | | | X | X | X | |
| ANR28 | 7.5 | X | | | X | X | X | |
| ANX11 | 1.6 | X | X | X | X | X | | |
| ANXA7 | 2.1 | X | | | X | X | X | |
| AP3S1 | 3.5 | X | X | | X | | X | |
| ARAF | 2.5 | X | | | X | | | |
| ARP10 | 3.1 | X | X | | X | X | | |
| AT1B1 | 2.9 | X | X | | X | | | |
| ATF6A | 5.3 | X | | | X | X | X | |
| ATG3 | 2.4 | X | | | | | | |
| ATG7 | 2.7 | X | X | | | | | X |
| ATG9A | 5.0 | X | | | X | | X | X |
| ATX2L | 2.2 | X | | | X | X | | |
| AVEN | 6.9 | X | | | | | | |
| BABA1 | 2.1 | X | | | X | X | | |
| BACH1 | 5.4 | X | | | X | X | | |
| BAG3 | 1.3 | X | X | | | | | |
| BAG6 | 2.7 | X | | | X | X | | |
| BCAR1 | 7.3 | X | X | | X | X | | |
| BCAR3 | 7.9 | X | | | X | | | |
| BICD2 | 8.5 | X | X | | X | X | X | |
| BIRC6 | 3.1 | X | X | X | X | | X | |
| BLM | 1.7 | X | | X | X | X | | |
| BRAT1 | 2.8 | X | | | | X | | |
| BUB1B | 2.6 | X | X | X | X | X | | |
| CACO2 | 7.3 | X | X | | X | X | | X |
| CAN1 | 2.9 | X | | | X | X | | |
| CAPZB | 1.8 | X | X | | X | | | |
| CCD50 | 2.8 | X | X | | | | | |
| CCDC6 | 1.5 | X | X | | | | | |
| CDC23 | 4.6 | X | | | X | X | | |
| CIP2A | 2.7 | X | | | | | | |
| CLAP2 | 2.2 | X | X | X | X | | X | |
| CLIP1 | 2.2 | X | X | X | X | X | | |
| CLU | 1.5 | X | | | | | | |
| CND1 | 1.5 | X | | X | X | X | | |
| CND2 | 1.4 | X | | | X | X | | |
| CNN2 | 1.4 | X | X | | | | | |
| CNO10 | 3.3 | X | | | X | X | | |
| CNOT1 | 3.2 | X | | | X | X | | |
| CNOT3 | 4.2 | X | | | X | X | | |
| COG2 | 2.3 | X | | | X | | X | |
| COG4 | 7.2 | X | X | | X | | X | |
| COG5 | 8.2 | X | | | X | X | X | |
| COG6 | 6.4 | X | | | X | | X | |
| CP1B1 | 3.6 | X | | | | | | |
| CSDE1 | 1.1 | X | | | X | | X | |
| CSN3 | 1.9 | X | | | X | X | | |
| CSN8 | 3.5 | X | | | X | X | | |
| CTU2 | 6.7 | X | | | X | | | |
| CYFP1 | 1.6 | X | X | | | | | |
| CYTSB | 6.0 | X | | | | | | |
| DBNL | 1.9 | X | X | | X | | X | |
| DC1I2 | 1.6 | X | X | X | | | | |
| DC1L1 | 1.3 | X | X | X | X | X | | |
| DCTN1 | 1.6 | X | X | X | X | X | | |
| DCTN2 | 1.9 | X | X | X | | | | |
| DCTN3 | 4.5 | X | X | X | | | | |
| DCTN4 | 3.1 | X | X | X | X | X | | |
| DDB1 | 2.3 | X | | | X | X | | |
| DEOC | 6.9 | X | | | X | X | | |
| DESP | 2.0 | X | X | | X | X | | |
| DIAP1 | 1.8 | X | X | X | | | | |
| DICER | 4.3 | X | X | | X | X | | |
| DJC13 | 2.2 | X | | | X | | | |
| DLG1 | 5.4 | X | X | | X | X | X | |
| DLGP5 | 4.6 | X | X | X | X | X | | |
| DNMBP | 5.8 | X | X | | | | X | |
| DTL | 5.6 | X | X | X | X | X | | |
| EEA1 | 2.9 | X | | | X | | | |
| EGFR | 1.5 | X | X | | X | X | X | |
| EH1L1 | 6.1 | X | | | X | | | |
| EIF2D | 2.1 | X | | | X | X | | |
| EIF3A | 1.6 | X | X | | X | X | | |
| EIF3B | 1.9 | X | | | X | X | | |
| EIF3C | 1.2 | X | | | X | | | |
| EIF3G | 1.7 | X | | | X | X | | |
| EIF3H | 1.5 | X | | | X | | | |
| EIF3I | 1.2 | X | | | X | | | |
| ELP1 | 2.5 | X | | | X | X | | |
| ELP3 | 1.2 | X | | | X | X | | |
| ELP5 | 6.5 | X | | | X | X | | |
| EP15R | 2.9 | X | | | X | X | | |
| EPIPL | 1.9 | X | X | | X | X | | |
| ERF3A | 1.3 | X | | | | | | |
| ES8L2 | 2.3 | X | | | | | | |
| ESYT1 | 2.8 | X | | | X | | X | |
| F120A | 1.3 | X | | | X | X | | |
| F175B | 3.3 | X | X | X | X | X | | |
| FADS1 | 2.3 | X | | | X | X | X | |
| FANCI | 1.5 | X | | | X | X | | |
| FAS | 1.1 | X | | | X | | X | |
| FKB15 | 8.3 | X | X | | X | | | |
| FKBP8 | 1.2 | X | | | X | | X | |
| FLII | 2.8 | X | X | | | X | | |
| FLNA | 1.6 | X | X | | X | X | | |
| FLNB | 2.3 | X | X | | | | | |
| FND3A | 2.0 | X | | | X | | X | |
| FOCAD | 5.8 | | X | | | | | |
| FOLC | 7.2 | X | | | | | | |
| FRPD3 | 4.6 | X | X | | | | | |
| FYCO1 | 2.1 | X | | | X | | | X |
| GAGE1 | 3.3 | X | | | | | | |
| GAPD1 | 2.6 | X | | | X | | | |
| GBF1 | 5.6 | X | | | X | | X | |
| GCC2 | 3.7 | X | | | X | | X | |
| GCP2 | 2.3 | X | X | X | X | X | | |
| GCR | 1.7 | X | X | X | X | X | | |
| GEPH | 5.9 | X | X | | | | | |
| GFPT2 | 2.8 | X | | | | | | |
| GID8 | 8.5 | X | | | X | X | | |
| GLCNE | 1.1 | X | | | | | | |

| | log2fold | cytoplasm/ cytosol | cyto-skeleton /cell junction | centro-some/ spindle | endosome/ trafficking | nucleus | endoplasmic reticulum/ golgi apparatus | pre-autophagosome/ autophagosome |
|---|---|---|---|---|---|---|---|---|
| GOGA2 | 2.4 | X | X | X | X | X | X | |
| GOGA3 | 4.3 | X | X | | X | X | X | |
| GOGA4 | 6.7 | X | | | X | X | X | |
| GOGB1 | 9.7 | X | | | | | X | |
| GOPC | 6.0 | X | X | | X | | X | |
| GRAM3 | 6.2 | X | X | | | | | |
| GSHR | 7.0 | X | | | X | | | |
| HAUS1 | 5.7 | X | X | | | X | | |
| HAUS6 | 7.5 | X | X | X | X | X | | |
| HAUS7 | 4.9 | X | X | | X | X | | |
| HBS1L | 6.2 | X | | | | | | |
| HCFC1 | 2.6 | X | X | | X | X | | |
| HEAT3 | 2.0 | X | | | | | | |
| HECD1 | 4.2 | X | | | | | | |
| HECD3 | 2.9 | X | | | | | | |
| HERC1 | 5.6 | X | | | X | | X | |
| HERC4 | 3.0 | X | | | X | X | | |
| HGS | 3.2 | X | | | X | | | |
| HN1L | 1.2 | X | | | X | X | | |
| HNRPL | 1.9 | X | | | X | X | | |
| HPRT | 2.0 | X | | | | | | |
| HS74L | 1.6 | X | | | X | X | | |
| HUWE1 | 3.8 | X | | | X | X | | |
| HXK2 | 1.2 | X | | | X | | | |
| ICAL | 2.2 | X | | | X | | | |
| ICLN | 2.2 | X | X | | X | X | | |
| IF4G1 | 1.1 | X | | | X | X | | |
| IFIT3 | 7.6 | X | | | X | | | |
| IMA5 | 3.7 | X | | | X | X | | |
| INF2 | 2.3 | X | | | | | | |
| IPO11 | 2.3 | X | | | X | X | | |
| IPO4 | 1.8 | X | | | X | X | | |
| IPO9 | 1.4 | X | | | X | X | | |
| IRAK1 | 3.3 | X | | | X | X | | |
| IRS2 | 2.7 | X | | | | | | |
| ITPR1 | 3.4 | X | | | X | X | X | |
| ITPR3 | 3.9 | X | | | X | X | X | |
| JIP4 | 3.2 | X | X | | X | | | |
| KEAP1 | 5.3 | X | X | | X | X | | |
| KI13A | 6.0 | X | X | | X | | X | |
| KIF11 | 2.3 | X | X | X | | | | |
| KIF14 | 3.8 | X | X | X | X | X | | |
| KIF15 | 5.4 | X | X | X | X | | | |
| KINH | 2.6 | X | X | | X | | | |
| KITH | 3.6 | X | | | | | | |
| KLC1 | 3.2 | X | X | | X | | | |
| KLC2 | 3.2 | X | X | | | | | |
| KPB2 | 8.8 | X | | | X | | | |
| KPBB | 4.7 | X | | | | | | |
| KPCI | 3.5 | X | X | | X | X | X | |
| LAMB3 | 3.3 | X | X | | | | | |
| LIMC1 | 2.5 | X | X | | | | | |
| LR16A | 1.6 | X | X | | X | X | | |
| LRBA | 5.9 | X | | | | | X | |
| LRCH3 | 4.0 | X | | | | | | |
| LRRF1 | 3.3 | X | X | | X | X | | |
| MACF1 | 3.1 | X | X | | | | | |
| MADD | 4.6 | X | | | | | | |
| MAEA | 3.2 | X | X | X | X | X | | |
| MAGD2 | 1.3 | X | | | | X | | |
| MALT1 | 8.3 | X | | | X | X | | |
| MAP4 | 1.9 | X | X | X | X | | | |
| MARC1 | 1.7 | X | | | X | | | |
| MBOA7 | 4.2 | X | | | X | | X | |
| MCMBP | 1.4 | X | | | X | X | | |
| MEP50 | 1.8 | X | | | X | X | X | |
| METH | 3.2 | X | | | | | | |
| MIA3 | 4.7 | X | | | X | | X | |
| MKLN1 | 7.8 | X | X | | | | | |
| ML12A | 1.6 | X | X | | | | | |
| MLP3B | 1.4 | X | X | | X | | | X |
| MON2 | 6.0 | X | | | X | | | |
| MTNB | 2.1 | X | | | | | | |
| MTOR | 3.5 | X | | | X | X | X | |
| MTSSL | 6.1 | X | X | | | | | |
| MYH9 | 1.3 | X | X | X | X | X | | |
| MYOF | 1.6 | X | | | X | X | | |
| MYPN | 3.1 | X | X | | | | | |
| MYPT1 | 2.7 | X | X | | X | X | | |
| NCDN | 8.2 | X | X | | | | | |
| NCKP1 | 3.2 | X | X | | | | | |
| NCOR1 | 5.3 | X | X | X | X | X | | |
| NDE1 | 5.8 | X | X | X | | | | |
| NEK9 | 4.3 | X | X | X | X | X | | |
| NFKB1 | 6.0 | X | | | X | X | | |
| NFKB2 | 2.7 | X | | | X | X | | |
| NP1L4 | 1.6 | X | | | X | X | | |
| NRBP | 2.3 | X | X | | X | X | | |
| NT5D1 | 2.5 | X | | | | X | | |
| NTKL | 2.6 | X | | | X | X | X | |
| NU133 | 2.7 | X | X | X | X | X | | |
| NU155 | 1.1 | X | | | X | X | | |
| NU214 | 2.3 | X | X | | X | X | | |
| NUBP2 | 1.1 | X | X | X | X | X | | |
| NUDC1 | 1.9 | X | | | X | X | | |
| NUF2 | 5.7 | X | X | X | X | | | |
| NUP50 | 4.2 | X | | | X | X | | |
| NUP88 | 2.1 | X | | | X | X | | |
| OGA | 2.8 | X | X | | X | X | | |
| OGFR | 2.7 | X | | | X | X | | |
| OGT1 | 3.0 | X | | | X | X | | |
| OSB11 | 3.4 | X | | | X | X | X | |
| OSBP1 | 3.4 | X | | | X | X | X | |
| OXA1L | 3.6 | X | | | X | | | |
| PA24A | 1.3 | X | | | X | | X | |
| PAB4L | 2.7 | X | | | | X | X | |
| PABP4 | 1.1 | X | | | X | X | | |
| PACN2 | 1.9 | X | X | | X | X | | |
| PAGE1 | 2.2 | X | | | | | | |
| PAIP1 | 2.9 | X | | | | | | |
| PALLD | 1.8 | X | X | | X | X | | |
| PANK4 | 2.2 | X | | | | | | |
| PAR12 | 3.0 | X | | | X | X | | |
| PARP4 | 3.8 | X | X | X | X | X | | |
| PAWR | 2.3 | X | X | | X | X | | |
| PCY1A | 2.5 | X | | | | X | | |
| PDC6I | 2.5 | X | X | | X | | X | |
| PDE3A | 3.2 | X | | | | | | |
| PDXD1 | 1.9 | X | | | | | X | |
| PHAX | 2.9 | X | X | X | X | X | | |
| PI3R4 | 1.4 | X | X | | X | X | | X |
| PJA2 | 4.7 | X | X | | X | | X | |
| PKN2 | 1.1 | X | X | X | X | X | | |
| PLEC | 1.4 | X | X | | | | | |
| PLIN3 | 1.5 | X | | | X | X | X | |
| PLIN4 | 2.8 | X | | | X | | | |
| PLK1 | 7.6 | X | X | X | X | X | | |
| PLXB2 | 2.9 | X | | | X | | | |
| PP4R1 | 3.6 | X | | | X | | | |
| PP6R3 | 3.9 | X | | | X | X | X | |
| PRC2C | 2.7 | X | | | | | | |
| PRDBP | 4.1 | X | | | | | | |
| PRS4 | 1.1 | X | | | X | X | | |
| PRS7 | 1.0 | X | | | X | X | | |
| PSA2 | 2.0 | X | | | X | X | | |
| PSA6 | 1.5 | X | X | | X | X | | |

| | log2fold | cytoplasm/ cytosol | cyto-skeleton /cell junction | centro-some/ spindle | endosome/ trafficking | nucleus | endoplasmic reticulum/ golgi apparatus | pre-autophagosome/ autophagosome |
|---|---|---|---|---|---|---|---|---|
| PSB1 | 1.5 | X | | | X | X | | |
| PSB5 | 1.7 | X | X | X | X | X | | |
| PSD12 | 1.5 | X | | | X | X | | |
| PSDE | 1.2 | X | | | X | X | | |
| PSME4 | 5.6 | X | | | X | X | | |
| PSMG2 | 2.7 | X | | | X | X | | |
| PTN11 | 1.6 | X | | | X | X | | |
| PTN12 | 3.6 | X | X | | X | X | | |
| PTN23 | 3.8 | X | X | | X | X | | |
| PUR6 | 1.8 | X | | | | | | |
| R3HD1 | 2.5 | X | | | | X | | |
| RABE1 | 7.0 | X | | | X | | | |
| RABE2 | 6.4 | X | | | X | | | |
| RANB9 | 4.6 | X | X | | X | X | | |
| RB3GP | 4.6 | X | | | X | | X | |
| RBGPR | 2.2 | X | | | X | | X | |
| RCD1 | 3.2 | X | | | X | X | | |
| RETST | 3.2 | X | | | X | X | X | |
| RIN1 | 4.4 | X | X | | | | | |
| RMD3 | 2.9 | X | X | X | | | | |
| ROCK1 | 6.5 | X | X | X | X | | X | |
| ROCK2 | 3.6 | X | X | X | X | X | | |
| RPB1 | 3.6 | X | | | X | X | | |
| RPB2 | 1.8 | X | | X | X | X | | |
| RPP30 | 1.5 | X | | | X | X | | |
| RS30 | 1.7 | X | | | X | X | | |
| RSU1 | 1.3 | X | X | | | | | |
| S23IP | 3.9 | X | | | X | | X | |
| S38A1 | 2.6 | X | | | | | | |
| S38A2 | 1.3 | X | | | | | | |
| SAC1 | 2.1 | X | | | X | | X | |
| SAS6 | 8.1 | X | X | X | | | | |
| SC23A | 2.5 | X | | | X | | X | |
| SC23B | 2.5 | X | | | X | | X | |
| SC24A | 3.9 | X | | | X | | X | |
| SC24B | 4.8 | X | | | X | | X | |
| SC24C | 2.4 | X | | | X | | X | |
| SC24D | 4.5 | X | | | X | | X | |
| SC31A | 2.1 | X | | | X | | X | |
| SCRIB | 2.7 | X | X | | X | X | | |
| SEPT6 | 2.0 | X | X | X | | | | |
| SEPT9 | 1.3 | X | X | | | | | |
| SH24A | 3.1 | X | | | | | | |
| SH3G1 | 1.5 | X | X | | X | | | |
| SHCBP | 3.5 | X | X | X | | | | |
| SHOT1 | 3.6 | X | X | | | | | |
| SMG9 | 4.1 | X | | | | | | |
| SNG2 | 3.6 | X | | | X | | | |
| SPAT5 | 2.9 | X | | | X | | | |
| SPCS3 | 1.2 | X | | | X | | X | |
| SPD2B | 2.7 | X | X | | | | | |
| SPG20 | 2.4 | X | | | X | | | |
| SPTN1 | 4.2 | X | X | | X | X | | |
| SQSTM | 3.6 | X | | | X | X | | X |
| SRC8 | 1.5 | X | | X | X | | X | |
| SRRM2 | 2.8 | X | | | X | X | | |
| SSA27 | 2.3 | | X | | | | | |
| STA5A | 2.2 | X | | | X | X | | |
| STAM1 | 3.0 | X | | | X | | | |
| STAM2 | 8.2 | X | | | X | X | | |
| STAT3 | 1.1 | X | | | X | X | | |
| STK24 | 3.3 | X | | | X | X | | |
| STRN | 3.4 | X | | | | | | |
| STRN4 | 6.3 | X | X | | X | | | |
| STX4 | 2.8 | X | X | | X | | X | |
| STXB3 | 2.0 | X | X | | X | | | |
| SYAP1 | 1.9 | X | | | X | X | X | |
| TAB2 | 5.7 | X | | | X | X | | |
| TACC2 | 1.7 | X | X | | X | X | | |
| TACC3 | 8.7 | X | X | X | | | | |
| TANC2 | 5.9 | X | X | | | | | |
| TAP1 | 1.6 | X | | | X | | X | |
| TAP2 | 1.5 | X | | | X | | X | |
| TARB1 | 4.3 | X | | | X | X | | |
| TAXB1 | 7.8 | X | | | | | | |
| TB182 | 3.0 | X | X | X | X | X | | |
| TBC15 | 2.6 | X | | | | | | |
| TBCD4 | 7.8 | X | | | X | | | |
| TBCD5 | 5.1 | X | | | X | | | X |
| TBD2A | 6.5 | X | | | X | | | |
| TBL3 | 2.9 | X | | | X | X | | |
| TELO2 | 3.0 | X | | X | X | X | | |
| TENS3 | 3.2 | X | X | | | | | |
| TENS4 | 7.4 | X | X | | | | | |
| TERA | 1.6 | X | | | X | X | | |
| TF3C5 | 2.0 | X | | | X | X | | |
| TF65 | 3.1 | X | | | X | X | | |
| TFG | 5.7 | X | | | X | | X | |
| TFP11 | 2.6 | X | | X | X | X | | |
| THUM3 | 1.9 | X | | | X | X | | |
| TIM21 | 2.6 | X | | | X | | | |
| TLN2 | 2.7 | X | X | | | | | |
| TMF1 | 7.7 | X | | | X | X | X | |
| TMOD3 | 1.6 | X | X | | | | | |
| TMX3 | 5.2 | X | | | X | | | |
| TNIP1 | 4.6 | X | | | X | X | | |
| TPM2 | 2.8 | X | X | | | | | |
| TPPC8 | 1.9 | X | X | | X | | X | |
| TRIP6 | 1.3 | X | X | | X | X | | |
| TRM1 | 1.1 | X | | | X | X | | |
| TRXR1 | 2.9 | X | | | X | X | | |
| TS101 | 4.2 | X | | | X | X | | |
| TTC1 | 1.6 | X | | | X | | | |
| TTC27 | 2.7 | | | X | | | | |
| TXLNA | 1.8 | X | | | | | | |
| TXLNG | 2.3 | X | | | X | X | | |
| UBA6 | 2.3 | X | | | | | | |
| UBAP2 | 2.7 | X | | | X | X | | |
| UBE2O | 3.7 | X | | | X | X | | |
| UBE3A | 3.1 | X | | | X | | | |
| UBE4A | 4.4 | X | | | X | X | | |
| UBP19 | 9.0 | X | | | X | | X | |
| UBP24 | 5.8 | X | | | | | | |
| UBP47 | 5.1 | X | | | X | X | | |
| UBR4 | 2.0 | X | X | | X | X | | |
| UCKL1 | 6.7 | X | | | | | | |
| UGPA | 1.6 | X | | | | | | |
| UN45A | 1.2 | X | | | X | X | | |
| URGCP | 3.6 | X | | | | | | |
| USP9X | 2.9 | X | X | | | | | |
| VASP | 1.5 | X | X | | | | | |
| VINC | 1.9 | X | X | | | | | |
| VP13C | 3.7 | X | | | X | | | |
| WBP2 | 5.6 | X | | | X | X | | |
| WDR6 | 3.7 | X | | | X | X | | |
| WDR81 | 2.5 | X | X | | X | X | | |
| WNK1 | 2.7 | X | | | | | | |
| YTDC2 | 3.0 | X | | | X | X | | |
| YTHD3 | 3.6 | X | | | | | | |
| ZBED1 | 3.4 | X | X | | X | X | | |
| ZFY16 | 7.8 | X | | | X | | | |
| ZN598 | 4.9 | | | | | | X | |
| ZNT1 | 2.8 | X | | | X | X | X | |
| ZPR1 | 7.6 | X | X | | X | X | | |
| ZW10 | 7.5 | X | X | X | | | | |

### Table S3 (related to Figure 4B)

#### Distribution of the 407 hits in pathways (DAVID)

##### AUTOPHAGY

| | |
|---|---|
| A16L1 | 3.0 |
| AAKG1 | 1.9 |
| ATG3 | 2.4 |
| ATG7 | 2.7 |
| ATG9A | 5.0 |
| CACO2 | 7.3 |
| FYCO1 | 2.1 |
| MLP3B | 1.4 |
| MTOR | 3.5 |
| PI3R4 | 1.4 |
| SQSTM | 3.6 |
| WDR6 | 3.7 |
| WDR81 | 2.5 |

##### FOCAL ADHESION-ACTIN CYTO-SKELETON -ECM

| | |
|---|---|
| ABL2 | 4.3 |
| ACTN1 | 3.4 |
| ACTN4 | 4.5 |
| ANKY2 | 3.6 |
| ANLN | 2.6 |
| ANR28 | 7.5 |
| ARP10 | 3.1 |
| BCAR1 | 7.3 |
| CAN1 | 2.9 |
| CAPZB | 1.8 |
| CCDC6 | 1.5 |
| CNN2 | 1.4 |
| CYFP1 | 1.6 |
| CYTSB | 6.0 |
| DBNL | 1.9 |
| DIAP1 | 1.8 |
| EPIPL | 1.9 |
| ES8L2 | 2.3 |
| FKB15 | 8.3 |
| FLII | 2.8 |
| FLNA | 1.6 |
| FLNB | 2.3 |
| FND3A | 2.0 |
| FOCAD | 5.8 |
| GEPH | 5.9 |
| GRAM3 | 6.2 |
| HAUS6 | 7.5 |
| HN1L | 1.2 |
| ICAL | 2.2 |
| INF2 | 2.3 |
| KIF11 | 2.3 |
| KIF14 | 3.8 |
| KPB2 | 8.8 |
| LAMB3 | 3.3 |
| LIMC1 | 2.5 |
| LR16A | 1.6 |
| LRCH3 | 4.0 |
| MACF1 | 3.1 |
| MKLN1 | 7.8 |
| ML12A | 1.6 |
| MTSSL | 6.1 |
| MYPN | 3.1 |
| MYPT1 | 2.7 |
| NCKP1 | 3.2 |
| PALLD | 1.8 |
| PLEC | 1.4 |
| PLXB2 | 2.9 |
| PTN11 | 1.6 |
| ROCK1 | 6.5 |
| ROCK2 | 3.6 |
| SEPT6 | 2.0 |
| SEPT9 | 1.3 |
| SHOT1 | 3.6 |
| SPD2B | 2.7 |
| SPTN1 | 4.2 |
| SRC8 | 1.5 |
| STK24 | 3.3 |
| TANC2 | 5.9 |
| TBD2A | 6.5 |
| TENS3 | 3.2 |
| TENS4 | 7.4 |
| TLN2 | 2.7 |
| TMOD3 | 1.6 |
| TPM2 | 2.8 |
| TRIP6 | 1.3 |
| TRXR1 | 2.9 |
| VASP | 1.5 |
| VINC | 1.9 |

##### MICROTUBULES

| | |
|---|---|
| BICD2 | 8.5 |
| CLAP2 | 2.2 |
| CLIP1 | 2.2 |
| DCTN3 | 4.5 |
| FKB15 | 8.3 |
| GCP2 | 2.3 |
| GEPH | 5.9 |
| GRAM3 | 6.2 |
| HAUS1 | 5.7 |
| HAUS6 | 7.5 |
| HAUS7 | 4.9 |
| HN1L | 1.2 |
| KI13A | 6.0 |
| KIF11 | 2.3 |
| KIF14 | 3.8 |
| KIF15 | 5.4 |
| MACF1 | 3.1 |
| MAP4 | 1.9 |
| NUF2 | 5.7 |
| RANB9 | 4.6 |
| RMD3 | 2.9 |
| SAS6 | 8.1 |
| TACC2 | 1.7 |
| ZW10 | 7.5 |

##### SIGNALLING

| | |
|---|---|
| AAKG1 | 1.9 |
| ABI1 | 8.6 |
| AKAP1 | 7.6 |
| ANM1 | 1.4 |
| ANX11 | 1.6 |
| ANXA7 | 2.1 |
| ARAF | 2.5 |
| BCAR3 | 7.9 |
| BRAT1 | 2.9 |
| CAN1 | 2.9 |
| CCD50 | 2.8 |
| CIP2A | 2.7 |
| DLG1 | 5.4 |
| EGFR | 1.5 |
| F120A | 1.3 |
| GEPH | 5.9 |
| IRS2 | 2.7 |
| JIP4 | 3.2 |
| KLC1 | 3.2 |
| KLC2 | 3.2 |
| KPB2 | 8.8 |
| KPBB | 4.7 |
| KPCI | 3.5 |
| MADD | 4.6 |
| ML12A | 1.6 |
| MTOR | 3.5 |
| MYPT1 | 2.7 |
| NCDN | 8.2 |
| PDE3A | 3.2 |
| PKN2 | 1.1 |
| PLK1 | 7.6 |
| PLXB2 | 2.9 |
| PP4R1 | 3.6 |
| PP6R3 | 3.9 |
| PRDBP | 4.1 |
| PTN11 | 1.6 |
| PTN12 | 3.6 |
| PTN23 | 3.8 |
| RANB9 | 4.6 |
| RIN1 | 4.4 |
| RSU1 | 1.3 |
| SH24A | 3.1 |
| SHCBP | 3.5 |
| STA5A | 2.2 |
| STAT3 | 1.1 |
| STK24 | 3.3 |
| STRN | 3.4 |
| TAB2 | 5.7 |
| TBCD4 | 7.8 |
| TF65 | 3.1 |
| TRIP6 | 1.3 |
| TRXR1 | 2.9 |
| VASP | 1.5 |
| WNK1 | 2.7 |
| ZFY16 | 7.8 |
| ZPR1 | 7.6 |

##### INFLAMMATION

| | |
|---|---|
| AIMP1 | 1.3 |
| GCR | 1.7 |
| HBS1L | 6.2 |
| HCFC1 | 2.6 |
| IFIT3 | 7.6 |
| IRAK1 | 3.3 |
| MALT1 | 8.3 |
| NFKB1 | 6.0 |
| NFKB2 | 2.7 |
| PA24A | 1.3 |
| PRS4 | 1.1 |
| STAT3 | 1.1 |
| TAB2 | 5.7 |
| TF65 | 3.1 |
| TNIP1 | 4.6 |
| TRIP6 | 1.3 |
| TRXR1 | 2.9 |

##### MEMBRANE TRANSPORTERS

| | |
|---|---|
| ABCD1 | 1.2 |
| AT1B1 | 2.9 |
| HEAT3 | 2.0 |
| HN1L | 1.2 |
| IMA5 | 3.7 |
| IPO11 | 2.3 |
| IPO4 | 1.8 |
| IPO9 | 1.4 |
| ITPR1 | 3.4 |
| ITPR3 | 3.9 |
| NU133 | 2.7 |
| NU155 | 1.1 |
| NU214 | 2.3 |
| NUP50 | 4.1 |
| NUP88 | 2.1 |
| S38A1 | 2.6 |
| S38A2 | 1.3 |
| TAP1 | 1.6 |
| TAP2 | 1.5 |
| TIM21 | 2.6 |
| ZNT1 | 2.8 |

##### PROTEIN MODIFICATION

| | |
|---|---|
| ALG1 | 2.3 |
| MEP50 | 1.8 |
| MTNB | 2.1 |
| OGT1 | 3.0 |

##### MITOCHONDRIA BIOGENESIS

| | |
|---|---|
| CLU | 1.5 |

##### RNA RELATED PROTEINS

| | | RNA transport | RNA degradation | RNA translation | RNA elongation | RNA splicing | tRNA modification | RNA Polymerase | RNA binding | micro RNA | mRNA surveillance pathway |
|---|---|---|---|---|---|---|---|---|---|---|---|
| AASD1 | 1.7 | | | | | | X | | | | |
| ABCE1 | 1.5 | | | | | | | | X | | |
| ACPH | 4.4 | | | | | | | | X | | |
| AHNK | 1.2 | | | | | X | | | | | |
| AIMP1 | 1.3 | | | | | | X | | | | |
| AK17A | 2.1 | | | | | X | | | | | |
| AKAP1 | 7.6 | | | | | | | | X | | |
| ANM5 | 2.6 | X | | | | | | | | | |
| ATX2L | 2.2 | | | | | | | | X | | |
| CNO10 | 3.3 | | X | | | | | | | | |
| CNOT1 | 3.2 | | X | | | | | | | | |
| CNOT3 | 4.2 | | X | | | | | | | | |
| CSDE1 | 1.1 | | | X | | | | | | | |
| CTU2 | 6.7 | | | | | | X | | | | |
| CYFP1 | 1.6 | X | | | | | | | | | |
| DICER | 4.3 | | | | | | | | | X | |
| EIF2D | 2.1 | | | X | | | | | | | |
| EIF3A | 1.6 | X | | | | | | | | | |
| EIF3B | 1.9 | X | | | | | | | | | |
| EIF3C | 1.2 | X | | | | | | | | | |
| EIF3G | 1.7 | X | | | | | | | | | |
| EIF3H | 1.5 | X | | | | | | | | | |
| EIF3I | 1.2 | X | | | | | | | | | |
| ELP1 | 2.5 | | | | X | | | | | | |
| ELP3 | 1.2 | | | | | | X | | | | |
| ELP5 | 6.5 | | | | X | | | | | | |
| ERF3A | 1.3 | | | X | | | | | | | |
| F120A | 1.3 | X | | | | | | | | | |
| HBS1L | 6.2 | | | | | | | | | | X |
| HNRPL | 1.9 | | | | | X | | | | | |
| ICLN | 2.2 | X | | | | | | | | | |
| IF4G1 | 1.1 | X | | | | | | | | | |
| NU133 | 2.7 | X | | | | | | | | | |
| NU155 | 1.1 | X | | | | | | | | | |
| NU214 | 2.3 | X | | | | | | | | | |
| NUP50 | 4.1 | X | | | | | | | | | |
| NUP88 | 2.1 | X | | | | | | | | | |
| PAB4L | 2.7 | X | X | | | | | | | | |
| PABP4 | 1.1 | X | X | | | | | | | | |
| PAIP1 | 2.9 | X | | | | | | | | | |
| PHAX | 2.9 | X | | | | | | | | | |
| PRC2C | 2.7 | | | X | | | | | | | |
| R3HD1 | 2.5 | | | | | | | | X | | |
| RCD1 | 3.2 | | X | | | | | | | | |
| RIN1 | 4.4 | | | | | | | | X | | |
| RPB1 | 3.6 | | | | | | | X | | | |
| RPB2 | 1.8 | | | | | | | X | | | |
| RPP30 | 1.5 | X | | | | | | | X | | |
| SMG9 | 4.1 | | X | | | | | | | | |
| SRRM2 | 2.8 | | | | | | | | X | | |
| TACC3 | 8.7 | X | | | | | | | | | |
| TARB1 | 4.3 | | | | | | X | | | | |
| TBL3 | 2.9 | | | | | | | | X | | |
| TFP11 | 2.6 | | | | | X | | | | | |
| THUM3 | 1.9 | | | | | | | | X | | |
| TRM1 | 1.1 | | | | | | X | | | | |
| TTC1 | 1.6 | | | | | | X | | | | |
| YTDC2 | 3.0 | | | | | | | | X | | |
| YTHD3 | 3.6 | | | X | | | | | | | |
| ZN598 | 4.9 | | | X | | | | | | | |

##### CELL-CELL JUNCTIONS

| | | Tight junction | Desmo-somes | Adherens Junctions | Cell polarity | Calcium binding protein |
|---|---|---|---|---|---|---|
| ACTN1 | 3.4 | X | | | | |
| ACTN4 | 4.5 | X | | | | |
| DESP | 2.0 | | X | | | |
| DLGP5 | 4.6 | | | X | | |
| KPCI | 3.5 | X | | | | |
| ML12A | 1.6 | X | | | | |
| MYH9 | 1.3 | X | | | | |
| SCRIB | 2.7 | | | | X | |
| SRC8 | 1.5 | X | | | | |
| STRN4 | 6.3 | | | | | X |
| TMX3 | 5.2 | | | | | X |

##### TRANSCRIPTION / DNA

| | | Transcription | DNA-Chromatin | DNA repair | Telomeres |
|---|---|---|---|---|---|
| BABA1 | 2.1 | | | X | |
| BACH1 | 5.4 | X | | | |
| BLM | 1.7 | | | X | |
| CND1 | 1.5 | | X | | |
| CND2 | 1.4 | | X | | |
| CNN2 | 1.4 | | X | | |
| DDB1 | 2.3 | | | X | |
| FANCI | 1.5 | | | X | |
| GCR | 1.7 | X | | | |
| LRRF1 | 3.3 | X | | | |
| MCMBP | 1.4 | | X | | |
| NCOR1 | 5.3 | X | | | |
| NP1L4 | 1.6 | X | X | | |
| NT5D1 | 2.5 | | X | | |
| PAR12 | 3.0 | X | | | |
| PARP4 | 3.8 | X | | | |
| SPAT5 | 2.9 | X | | | |
| TARB1 | 4.3 | X | | | |
| TB182 | 3.0 | X | | | |
| TELO2 | 3.0 | | | | X |
| TF3C5 | 2.0 | X | | | |
| TMF1 | 7.7 | X | | | |
| TRXR1 | 2.9 | X | | | |
| TXLNA | 1.8 | X | | | |
| TXLNG | 2.3 | X | | | |
| WBP2 | 5.6 | X | | | |
| ZBED1 | 3.4 | X | | | |

##### NUCLEAR ENVELOPPE

| | |
|---|---|
| ANKL2 | 4.1 |

##### CELL CYCLE / MITOSIS / CYTOKINESIS

| | | Cell cyle/ mi-tosis | Cytoki-nesis |
|---|---|---|---|
| ANLN | 2.6 | | X |
| ANX11 | 1.6 | | X |
| BUB1B | 2.6 | X | |
| DCTN3 | 4.5 | | X |
| DLGP5 | 4.6 | X | |
| HAUS1 | 5.7 | | X |
| HAUS6 | 7.5 | X | |
| HAUS7 | 4.9 | | X |
| HCFC1 | 2.6 | X | |
| KI13A | 6.0 | X | |
| MAGD2 | 1.3 | X | |
| ML12A | 1.6 | | X |
| NDE1 | 5.8 | | X |
| NEK9 | 4.3 | X | |
| NRBP | 2.3 | X | |
| NUF2 | 5.7 | X | |
| OGFR | 2.7 | X | |
| SEPT9 | 1.3 | | X |
| SPAT5 | 2.9 | X | |
| SSA27 | 2.3 | X | |

##### APOPTOSIS / SURVIVAL

| | |
|---|---|
| AVEN | 6.9 |
| BAG3 | 1.3 |
| BAG6 | 2.7 |
| CAN1 | 2.9 |
| FKBP8 | 1.2 |
| PAWR | 2.3 |
| TAXB1 | 7.8 |

##### ANTIGEN PROCESSING AND PRESENTATION OF EXOGENOUS PEPTIDE ANTIGEN VIA MHC CLASS I or II

| | | Class II | Class I |
|---|---|---|---|
| AT1B1 | 2.9 | X | |
| KLC1 | 3.2 | X | |
| KLC2 | 3.2 | X | |
| TAP1 | 1.6 | | X |
| TAP2 | 1.5 | | X |

##### PROLIFERATION

| | |
|---|---|
| UCKL1 | 6.7 |
| URGCP | 3.6 |
| ZPR1 | 7.6 |

##### UNKNOWN FUNCTION

| | |
|---|---|
| GAGE1 | 3.3 |
| NUDC1 | 1.9 |
| PAGE1 | 2.2 |
| TTC27 | 2.7 |

##### RIBOSOMAL PROTEIN

| | |
|---|---|
| RS30 | 1.7 |

##### METABOLISM

| | |
|---|---|
| A16A1 | 5.5 |
| AAKG1 | 1.9 |
| ABCD1 | 1.2 |
| ACLY | 1.8 |
| ALDOA | 1.2 |
| AMPD2 | 2.9 |
| BRAT1 | 2.9 |
| CP1B1 | 3.6 |
| DEOC | 6.9 |
| FADS1 | 2.3 |
| FAS | 1.1 |
| FOLC | 7.2 |
| GCR | 1.7 |
| GFPT2 | 2.8 |
| GLCNE | 1.1 |
| GSHR | 7.0 |
| HPRT | 2.0 |
| HXK2 | 1.2 |
| KITH | 3.6 |
| KPB2 | 8.8 |
| MARC1 | 1.7 |
| MBOA7 | 4.2 |
| METH | 3.2 |
| NUBP2 | 1.1 |
| OGA | 2.8 |
| OSB11 | 3.4 |
| OSBP1 | 3.4 |
| OXA1L | 3.6 |
| PA24A | 1.3 |
| PANK4 | 2.2 |
| PCY1A | 2.5 |
| PDXD1 | 1.9 |
| PLIN3 | 1.5 |
| PLIN4 | 2.8 |
| PUR6 | 1.8 |
| RETST | 3.2 |
| SAC1 | 2.1 |
| TERA | 1.6 |
| TIM21 | 2.6 |
| TRXR1 | 2.9 |
| UCKL1 | 6.7 |
| UGPA | 1.6 |
| WNK1 | 2.7 |

##### ENDOCYTOSIS- TRAFFICKING-UBIQUITINATION-PROTEASOME

| | | Endocytosis and membrane trafficking | Protein processing in endoplasmic reticulum | Ubiquitin mediated proteolysis | Protea-some | Chape-rone |
|---|---|---|---|---|---|---|
| ACAP2 | 4.2 | X | | | | |
| AP3S1 | 3.5 | X | | | | |
| ATF6A | 5.3 | | X | | | |
| BABA1 | 2.1 | | | X | | |
| BICD2 | 8.5 | X | | | | |
| BIRC6 | 3.1 | | | X | | |
| CAN1 | 2.9 | | X | | | |
| CAPZB | 1.8 | X | | | | |
| CDC23 | 4.6 | | | X | | |
| CLIP1 | 2.2 | X | | | | |
| COG2 | 2.3 | X | | | | |
| COG4 | 7.2 | X | | | | |
| COG5 | 8.2 | X | | | | |
| COG6 | 6.4 | X | | | | |
| CSN3 | 1.9 | X | | X | | |
| CSN8 | 3.5 | X | | X | | |
| DBNL | 1.9 | X | | | | |
| DC1I2 | 1.6 | X | | | | |
| DC1L1 | 1.3 | X | | | | |
| DCTN1 | 1.6 | X | | | | |
| DCTN2 | 1.9 | X | | | | |
| DCTN3 | 4.5 | X | | | | |
| DCTN4 | 3.1 | X | | | | |
| DDB1 | 2.3 | | | X | | |
| DJC13 | 2.2 | X | | | | |
| DNMBP | 5.8 | X | | | | |
| DTL | 5.6 | | | X | | |
| EEA1 | 2.9 | X | | | | |
| EH1L1 | 6.1 | X | | | | |
| EP15R | 2.9 | X | | | | |
| ESYT1 | 2.8 | X | | | | |
| F175B | 3.3 | | | X | | |
| FKB15 | 8.3 | X | | | | |
| FRPD3 | 4.6 | X | | | | |
| FYCO1 | 2.1 | X | | | | |
| GAPD1 | 2.6 | X | | | | |
| GBF1 | 5.6 | X | | | | |
| GCC2 | 3.7 | X | | | | |
| GID8 | 8.5 | | | X | | |
| GOGA2 | 2.4 | X | | | | |
| GOGA3 | 4.3 | X | | | | |
| GOGA4 | 6.7 | X | | | | |
| GOGB1 | 9.7 | X | | | | |
| GOPC | 6.0 | X | | | | |
| HECD1 | 4.2 | | | X | | |
| HECD3 | 2.9 | | | X | | |
| HERC1 | 5.6 | | | X | | |
| HERC4 | 3.0 | | | X | | |
| HGS | 3.2 | X | | | | |
| HS74L | 1.6 | | X | | | |
| HUWE1 | 3.8 | | | X | | |
| JIP4 | 3.2 | X | | | | |
| KEAP1 | 5.3 | | | X | | |
| KINH | 2.6 | X | | | | |
| KLC1 | 3.2 | | X | | | |
| KLC2 | 3.2 | | X | | | |
| KPCI | 3.5 | X | | | | |
| LRBA | 5.9 | X | | | | |
| MADD | 4.6 | X | | | | |
| MAEA | 3.2 | | | X | | |
| MIA3 | 4.7 | | X | | | |
| MKLN1 | 7.8 | X | | X | | |
| MON2 | 6.0 | X | | | | |
| MYOF | 1.6 | X | | | | |
| NDE1 | 5.8 | X | | | | |
| NTKL | 2.6 | | X | | | |
| PACN2 | 1.9 | X | | | | |
| PDC6I | 2.5 | X | | | | |
| PJA2 | 2.6 | | | X | | |
| PRS4 | 1.1 | | | | X | |
| PRS7 | 1.0 | | | | X | |
| PSA2 | 2.0 | | | | X | |
| PSA6 | 1.5 | | | | X | |
| PSB1 | 1.5 | | | | X | |
| PSB5 | 1.7 | | | | X | |
| PSD12 | 1.5 | | | | X | |
| PSDE | 1.2 | | | | X | |
| PSME4 | 5.6 | | | | X | |
| PSMG2 | 2.7 | | | | X | |
| PTN23 | 3.8 | X | | | | |
| RABE1 | 7.0 | X | | | | |
| RABE2 | 6.4 | X | | | | |
| RB3GP | 4.6 | X | | | | |
| RBGPR | 2.2 | X | | | | |
| RIN1 | 4.4 | X | | | | |
| S23IP | 3.9 | | X | | | |
| SC23A | 2.5 | | X | | | |
| SC23B | 2.5 | X | X | | | |
| SC24A | 3.9 | | X | | | |
| SC24B | 4.8 | | X | | | |
| SC24C | 2.4 | | X | | | |
| SC24D | 4.5 | | X | | | |
| SC31A | 2.1 | | X | | | |
| SEPT9 | 1.3 | X | | | | |
| SH3G1 | 1.5 | X | | | | |
| SNG2 | 3.6 | X | | | | |
| SPAT5 | 2.9 | X | | X | | |
| SPCS3 | 1.2 | | X | | | |
| SPG20 | 2.4 | X | | | | |
| STAM1 | 3.0 | X | | | | |
| STAM2 | 8.2 | X | | | | |
| STX4 | 2.8 | X | | | | |
| STXB3 | 2.0 | X | | | | |
| SYAP1 | 1.9 | X | | | | |
| TAP1 | 1.6 | X | | | | |
| TAP2 | 1.5 | X | | | | |
| TBC15 | 2.6 | X | | | | |
| TBCD4 | 7.8 | X | | | | |
| TBCD5 | 5.1 | X | | | | |
| TBD2A | 6.5 | X | | | | |
| TERA | 1.6 | X | X | X | | |
| TFG | 5.7 | X | | | | |
| TMF1 | 7.7 | X | | | | |
| TPPC8 | 1.9 | X | | | | |
| TS101 | 4.2 | X | | | | |
| TXLNA | 1.8 | X | | | | |
| TXLNG | 2.3 | X | | | | |
| UBA6 | 2.3 | | | X | | |
| UBAP2 | 2.7 | | | X | | |
| UBE2O | 3.7 | | | X | | |
| UBE3A | 3.1 | | | X | | |
| UBE4A | 4.4 | | | X | | |
| UBP19 | 9.0 | | | X | | |
| UBP24 | 5.8 | | | X | | |
| UBP47 | 5.1 | | | X | | |
| UBR4 | 2.0 | | | X | | |
| UN45A | 1.2 | | | | | X |
| USP9X | 2.9 | | | X | | |
| VP13C | 3.7 | X | | | | |
| WDR81 | 2.5 | X | | | | |
| ZFY16 | 7.8 | X | | | | |
| ZN598 | 4.9 | | | X | | |

### Figure S4 (related to Figure 4CD)

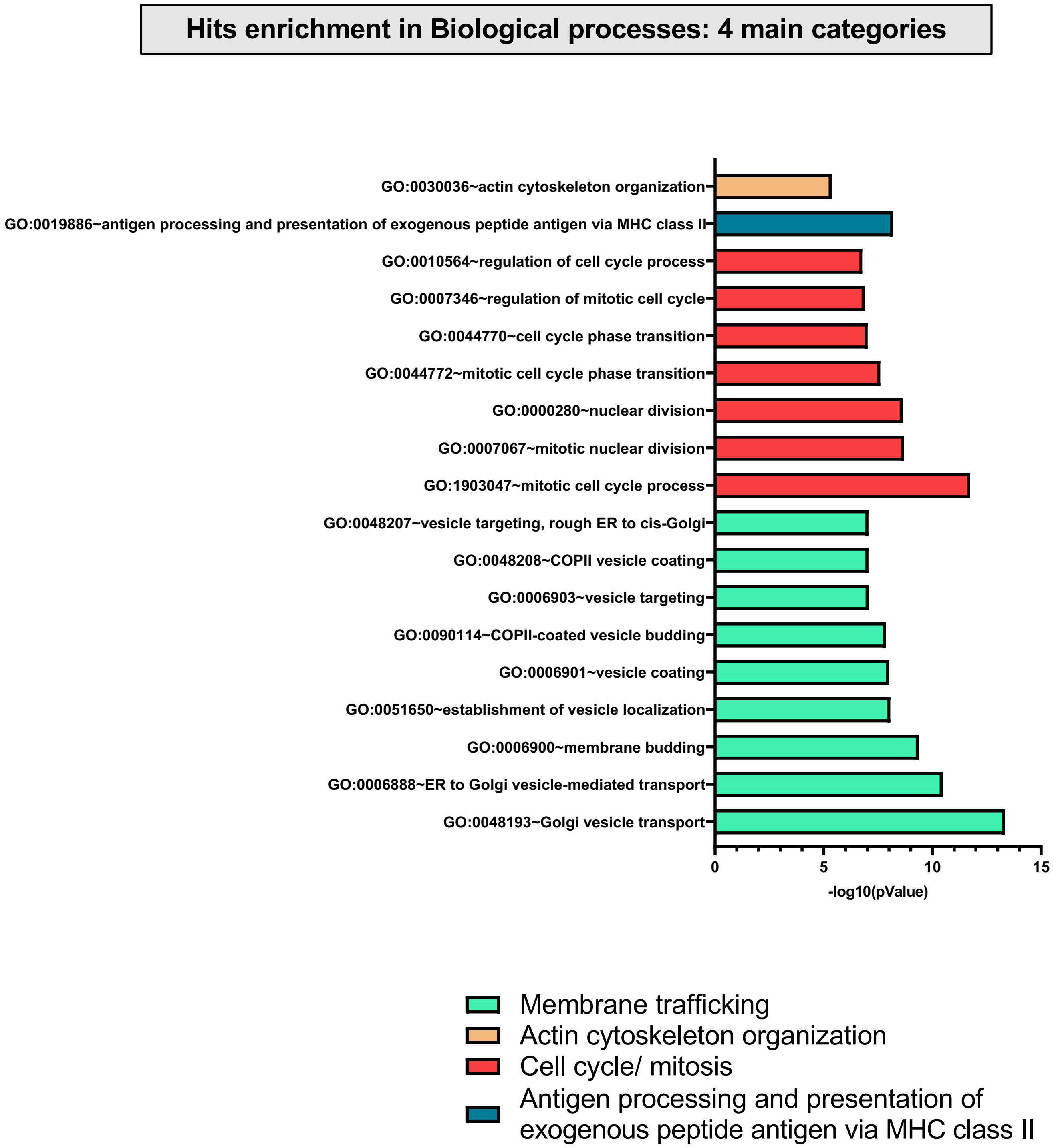

### Figure S5 (related to Figure 4CD)

#### Distribution of the hits enriched following GO "Biological processes" analysis

##### Previously reported to be related to autophagy

| Gene | | Membrane trafficking | Actin cytoskeleton organization | Cell cycle/ mitosis | Antigen processing and presentation of exogenous peptide antigen via MHC class II | Refs | LC3 asso-ciation |
|---|---|---|---|---|---|---|---|
| ACTN4 | 4.5 | X | X | | | Yee et al., 2018 | |
| ANM5 | 2.6 | | | X | | Chitiprolu et al., 2018 | |
| BACH1 | 5.4 | | | X | | Takada et al., 2015 | |
| BAG6 | 2.7 | | | X | | Ragimbeau et al., 2021 | X |
| BCAR1 | 7.3 | | X | | | Bisaro et al., 2015 | |
| BIRC6 | 3.1 | | | X | | Jia and Bonifacino, 2019 | X |
| CNOT1 | 3.2 | | | X | | Yamaguchi et al., 2018 | |
| CNOT3 | 4.2 | | | X | | Yamaguchi et al., 2018 | |
| DCTN1 | 1.6 | X | | X | X | Wang et al., 2020 | |
| DCTN2 | 1.9 | X | | X | X | Xu et al., 2014 | |
| DIAP1 | 1.8 | | X | | | Bargiela et al., 2015 | |
| EGFR | 1.5 | | | X | | Tan et al., 2016 | X |
| FANCI | 1.5 | | | X | | Sun et al., 2020 | |
| FYCO1 | 2.1 | X | | | | Pankiv & Johansen, 2010 | X |
| GCR | 1.7 | | | X | | Lee et al., 2018 | |
| GOGA2 | 2.4 | X | | X | | Chang et al., 2012 | |
| GOGA4 | 6.7 | X | | | | Sohda et al., 2015 | |
| IF4G1 | 1.1 | | | X | | Ramírez-Valle et al., 2008 | |
| KINH | 2.6 | X | | | | Chen and Yu, 2017 | |
| KPCI | 3.5 | X | X | | | Qu et al., 2016 | |
| MTOR | 3.5 | | X | | | Kim et al., 2011 | |
| MYH9 | 1.3 | | X | | | Ibrahim et al., 2017 | |
| NEK9 | 4.3 | | | X | | Shrestha et al., 2019 | X |
| PAWR | 2.3 | | X | | | Rah et al., 2015 | |
| PDC6I | 2.5 | X | | X | | Murrow et al., 2015 | |
| PI3R4 | 1.4 | | | X | | Birgisdottir et al., 2019 | X |
| PLK1 | 7.6 | | | X | | Ruf et al., 2017 | |
| RB3GP | 4.6 | X | | | | Spang et al., 2014 | X |
| ROCK1 | 6.5 | | X | | | Gurkar et al., 2013 | |
| ROCK2 | 3.6 | | X | X | | Shi et al., 2019 | |
| SC23A | 2.5 | X | | | X | Gan et al., 2017 | |
| SC24A | 3.9 | X | | | X | Jeong et al., 2018 | |
| SC24B | 4.8 | X | | | X | Jeong et al., 2018 | |
| SC24C | 2.4 | X | | | X | Cui et al., 2019 | |
| SC24D | 4.5 | X | | | X | Stadel et al., 2015 | |
| SC31A | 2.1 | X | | | X | Cleyrat et al., 2014 | |
| TFG | 5.7 | X | | | | Steinmetz et al., 2020 | |
| TS101 | 4.2 | X | | | | Morris et al, 2012 | |
| TXLNG | 2.3 | | | X | | Rah et al., 2015 | |
| UBP19 | 9 | | | X | | Jin et al., 2016 | |
| USP9X | 2.9 | | | X | | Grasso et al., 2011 | |
| VASP | 1.5 | | X | | | Kiss et al., 2019 | |
| VP13C | 3.7 | X | | | | Lesage et al., 2016 | |
| WDR81 | 2.5 | X | | | X | Liu et al., 2017 | X |

##### Not previously reported to be related to autophagy

| Gene | | Membrane trafficking | Actin cytoskeleton organization | Cell cycle/ mitosis | Antigen processing and presentation of exogenous peptide antigen via MHC class II |
|---|---|---|---|---|---|
| ABI1 | 8.6 | | X | | |
| ACTN1 | 3.4 | | X | | |
| ALDOA | 1.2 | | X | | |
| ANKL2 | 4.1 | | | X | |
| ANLN | 2.6 | | | X | |
| ANM1 | 1.4 | | | X | |
| ANR28 | 7.5 | X | | | |
| BABA1 | 2.1 | | | X | |
| BICD2 | 8.5 | X | | | |
| BUB1B | 2.6 | | | X | |
| CDC23 | 4.6 | | | X | |
| CLIP1 | 2.2 | | | X | |
| CND1 | 1.5 | | | X | |
| CND2 | 1.4 | | | X | |
| CNN2 | 1.4 | | X | | |
| CNO10 | 3.3 | | | X | |
| COG5 | 8.2 | X | | | |
| COG6 | 6.4 | X | | | |
| CYFP1 | 1.6 | | X | | |
| DC1I2 | 1.6 | X | | X | X |
| DC1L1 | 1.3 | | | X | |
| DCTN3 | 4.5 | X | | X | X |
| DCTN4 | 3.1 | X | | | X |
| DLG1 | 5.4 | | X | X | |
| DLGP5 | 4.6 | | | X | |
| ERF3A | 1.3 | | | X | |
| F175B | 3.3 | | | X | |
| FLNB | 2.3 | | X | | |
| GBF1 | 5.6 | X | | X | |
| GCC2 | 3.7 | X | | | |
| HAUS1 | 5.7 | | | X | |
| HAUS7 | 4.9 | | | X | |
| INF2 | 2.3 | | X | | |
| KIF11 | 2.3 | X | | X | X |
| KLC1 | 3.2 | X | | | X |
| KLC2 | 3.2 | X | | | X |
| MAEA | 3.2 | | | X | |
| MCMBP | 1.4 | | | X | |
| MIA3 | 4.7 | X | | | |
| MKLN1 | 7.8 | | X | | |
| MON2 | 6 | X | | | |
| NCKP1 | 3.2 | | X | | |
| NDE1 | 5.8 | X | | X | |
| NRBP | 2.3 | X | | | |

| Gene | | Membrane trafficking | Actin cytoskeleton organization | Cell cycle/ mitosis | Antigen processing and presentation of exogenous peptide antigen via MHC class II |
|---|---|---|---|---|---|
| NU155 | 1.1 | | | X | |
| NUP50 | 4.2 | | | X | |
| PACN2 | 1.9 | | X | | |
| PALLD | 1.8 | | X | | |
| PDE3A | 3.2 | | | X | |
| PKN2 | 1.1 | | | X | |
| PP6R3 | 3.9 | X | | | |
| PRDBP | 4.1 | | X | | |
| PSB5 | 1.7 | | | X | |
| PSD12 | 1.5 | | | X | |
| PSMG2 | 2.7 | | | X | |
| RCD1 | 3.2 | | | X | |
| S23IP | 3.9 | X | | | |
| SHOT1 | 3.6 | | X | | |
| SRC8 | 1.5 | | X | | |
| SSA27 | 2.3 | | | X | |
| STX4 | 2.8 | X | | | |
| TACC3 | 8.7 | | | X | |
| TB182 | 3 | | | X | |
| TPM2 | 2.8 | | X | | |
| TXLNG | 2.3 | | | X | |
| UBP47 | 5.1 | | | X | |

### Figure S6 (related to Figure 4E)

#### High confidence hits

**A**

| Previously reported to be autophagy- related | | | Reported association with LC3 |
|---|---|---|---|
| | Log2fold | References | |
| ABCE1 | 1.5 | Wu et al., 2018 | |
| ACLY | 1.8 | Mariño et al., 2014 | |
| ANM5 | 2.6 | Chitiprolu et al., 2018 | |
| BAG6 | 2.7 | Ragimbeau et al., 2021 | |
| BCAR1 | 7.3 | Bisaro et al., 2015 | |
| BIRC6 | 3.1 | Jia and Bonifacino, 2019 | x |
| DCTN2 | 1.9 | Xu et al., 2014 | |
| EEA1 | 2.9 | Wu et al., 2016 | x |
| ESYT1 | 2.8 | Pan et al., 2020 | |
| FLII | 2.8 | He et al., 2018 | |
| FLNA | 1.6 | Wang et al., 2017 | |
| GBF1 | 5.6 | Naydenov et al., 2012 | |
| HECD1 | 4.2 | Han et al., 2018 | |
| HUWE1 | 3.8 | Wan et al., 2018 | |
| ICAL | 2.2 | Santarelli et al., 2016 | |
| IRS2 | 2.7 | Sadagurski et al., 2011 | |
| KINH | 2.6 | Chen and Yu, 2017 | |
| KPCI | 3.5 | Qu et al., 2016 | |
| MYOF | 1.6 | Han et al., 2019 | |
| NEK9 | 4.3 | Shrestha et al., 2019 | x |
| OGA | 2.8 | Zhu et al., 2018 | |
| OGT1 | 3 | Guo et al., 2014 | |
| PDC6I | 2.5 | Murrow et al., 2015 | |
| PLIN3 | 1.5 | Kaushik et al., 2015 | |
| PLIN4 | 2.8 | Han et al., 2018 | |
| RBGPR | 2.2 | Spang et al., 2014 | |
| SC24B | 4.8 | Jeong et al., 2018 | |
| SC24C | 2.4 | Cui et al., 2020 | |
| TB182 | 3 | Wang et al., 2020 | |
| TF65 | 3.1 | Nopparat et al., 2017 | |
| TFG | 5.7 | Steinmetz et al., 2020 | |
| TS101 | 4.2 | Morris et al., 2012 | |
| UBR4 | 2 | Tasaki et al., 2013 | x |
| USP9X | 2.9 | Ma et al., 2018 | |
| WNK1 | 2.7 | Gallolu Kankanamalage et al., 2017 | |

| Not previously reported to be autophagy-related | |
|---|---|
| | Log2fold |
| AIMP1 | 1.3 |
| AMPD2 | 2.9 |
| ANM1 | 1.4 |
| CDC23 | 4.6 |
| CLIP1 | 2.2 |
| CND1 | 1.5 |
| DCTN4 | 3.1 |
| EIF3A | 1.6 |
| EIF3B | 1.9 |
| EIF3H | 1.5 |
| EIF3I | 1.2 |
| ELP1 | 2.5 |
| FKB15 | 8.3 |
| FLNB | 2.3 |
| GAPD1 | 2.6 |
| IFIT3 | 7.6 |
| INF2 | 2.3 |
| JIP4 | 3.2 |
| KIF11 | 2.3 |
| KIF15 | 5.4 |
| KLC1 | 3.2 |
| KLC2 | 3.2 |
| LIMC1 | 2.5 |
| LRRF1 | 3.3 |
| MAP4 | 1.9 |
| MKLN1 | 7.8 |
| MYPN | 3.1 |
| NCKP1 | 3.2 |
| NFKB2 | 2.7 |
| OSB11 | 3.4 |
| OSBP1 | 3.4 |
| OXA1L | 3.6 |
| PABP4 | 1.1 |
| PDE3A | 3.2 |
| PDXD1 | 1.9 |
| PSMG2 | 2.7 |
| PTN12 | 3.6 |
| RABE1 | 7 |
| S23IP | 3.9 |
| SCRIB | 2.7 |
| SHCBP | 3.5 |
| SHOT1 | 3.6 |
| SPD2B | 2.7 |
| SPG20 | 2.4 |
| SRC8 | 1.5 |
| STAM1 | 3 |
| TERA | 1.6 |
| TTC27 | 2.7 |
| UBE2O | 3.7 |
| UBP47 | 5.1 |
| UN45A | 1.2 |
| ZFY16 | 7.8 |
| ZPR1 | 7.6 |

**B**

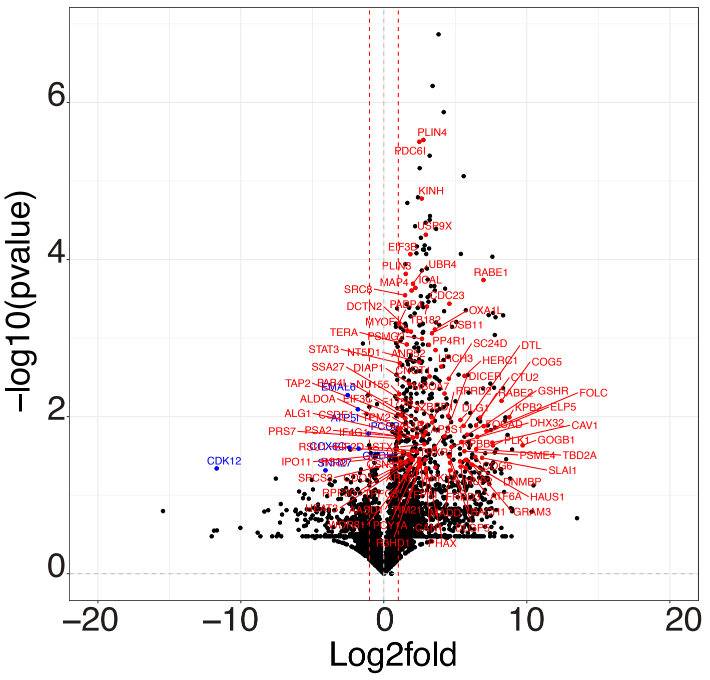

### Figure S7 (related to Figure 5A)

#### Comparison with four previous studies

##### Previously reported to be related to autophagy

| | log2fold | Le guerroué et al. | Dengjel et al. | Gao et al. | Mancias et al. | Reported association with LC3 |
|---|---|---|---|---|---|---|
| AAKG1 | 1.9 | | X | X | | |
| ABCE1 | 1.5 | X | X | | | |
| ACLY | 1.8 | X | X | | | |
| ACTN4 | 4.5 | X | X | | | |
| ANM5 | 2.6 | | X | | | |
| ANXA7 | 2.1 | | X | | | X |
| ATG7 | 2.7 | | | X | | X |
| ATG9A | 5.0 | | | X | X | |
| BAG3 | 1.3 | | X | | X | |
| CACO2 | 7.3 | | X | | X | X |
| CAPZB | 1.8 | X | X | | | |
| CLU | 1.5 | X | X | | X | X |
| CNOT1 | 3.2 | | X | | | |
| CP1B1 | 3.6 | | X | | | |
| DCTN1 | 1.6 | X | | | | |
| DDB1 | 2.3 | | X | | | |
| DJC13 | 2.2 | X | X | | | |
| EEA1 | 2.9 | | X | | | X |
| EGFR | 1.5 | X | | | | X |
| EIF3G | 1.7 | X | | | | |
| ESYT1 | 2.8 | X | | | | |
| FAS | 1.1 | X | X | | | |
| FKBP8 | 1.2 | | X | | | X |
| FLII | 2.8 | | X | | | |
| FLNA | 1.6 | X | X | | | |
| FYCO1 | 2.1 | | | | X | X |
| GBF1 | 5.6 | | X | | | |
| GOGA2 | 2.4 | | X | | | |
| GOPC | 6.0 | | | | X | |
| HGS | 3.2 | | X | | X | X |
| HPRT | 2.0 | | X | | | |
| HUWE1 | 3.8 | | X | | | |
| IF4G1 | 1.1 | X | X | | | |
| ITPR1 | 3.4 | | X | | | |
| ITPR3 | 3.9 | | X | | | |
| KEAP1 | 5.3 | | | | X | X |
| KINH | 2.6 | X | | | | |
| KPCI | 3.5 | | X | | | |
| MACF1 | 3.1 | | X | | | |
| MLP3B | 1.4 | | X | X | X | X |
| MYH9 | 1.3 | X | | | | |
| MYOF | 1.6 | X | | | | |
| PAWR | 2.3 | | X | | | |
| PDC6I | 2.5 | X | X | | X | |
| PLEC | 1.4 | X | X | | | |
| PSDE | 1.2 | | X | | | |
| RBGPR | 2.2 | | X | | X | |
| ROCK1 | 6.5 | | X | | | |
| ROCK2 | 3.6 | | X | | | |
| S38A1 | 2.6 | | X | | | |
| S38A2 | 1.3 | | X | | X | |
| SC23A | 2.5 | | X | | | |
| SC24C | 2.4 | X | X | | | |
| SC31A | 2.1 | X | X | | | |
| SQSTM | 3.6 | X | X | | X | X |
| STAM2 | 8.2 | | X | | X | |
| STK24 | 3.3 | | X | | | |
| TAXB1 | 7.8 | | X | | X | X |
| TBC15 | 2.6 | | | | X | X |
| TFG | 5.7 | | X | | | |
| TNIP1 | 4.6 | | | | X | |
| TRXR1 | 2.9 | X | | | | |
| TS101 | 4.2 | | X | | X | |
| UBR4 | 2.0 | | X | | | X |
| USP9X | 2.9 | | X | | | |
| VASP | 1.5 | | X | | | |
| VP13C | 3.7 | | | | X | |
| WBP2 | 5.6 | | | | X | |

##### Not previously reported to be related to autophagy

| | Log2fold | Le guerroué et al. | Dengjel et al. | Gao et al. | Mancias et al. |
|---|---|---|---|---|---|
| ABI1 | 8.6 | X | | | |
| ACTN1 | 3.4 | X | X | | |
| AHNK | 1.2 | | X | | |
| ALDOA | 1.2 | X | | | |
| ALG1 | 2.3 | | X | | |
| AT1B1 | 2.9 | X | X | | |
| CND2 | 1.4 | | X | | |
| CSDE1 | 1.1 | X | | | |
| CYFP1 | 1.6 | | X | | |
| DCTN3 | 4.5 | | | | X |
| DESP | 2.0 | X | X | | |
| DLG1 | 5.4 | | X | | |
| EIF3A | 1.6 | X | | | |
| EIF3B | 1.9 | X | | | |
| EIF3I | 1.2 | X | | | |
| EP15R | 2.9 | | X | | |
| EPIPL | 1.9 | X | | | |
| ERF3A | 1.3 | X | | | |
| F120A | 1.3 | | X | | |
| FKB15 | 8.3 | | | | X |
| FLNB | 2.3 | X | X | | |
| GCP2 | 2.3 | | | X | |
| GOGB1 | 9.7 | | X | | |
| GSHR | 7.0 | | X | | |
| HNRPL | 1.9 | X | | | |
| IPO4 | 1.8 | X | X | | |
| LIMC1 | 2.5 | | X | | |
| LR16A | 1.6 | | X | | |
| MAGD2 | 1.3 | | X | | |
| MAP4 | 1.9 | | X | | |
| MIA3 | 4.7 | X | X | | |
| ML12A | 1.6 | X | | | |
| MYPT1 | 2.7 | | | | X |
| NCKP1 | 3.2 | | X | | |
| NFKB2 | 2.7 | X | | | |
| NTKL | 2.6 | | | | X |
| OGFR | 2.7 | X | | | |
| OSBP1 | 3.4 | | | | X |
| OXA1L | 3.6 | X | | | |
| PACN2 | 1.9 | X | X | | |
| PAIP1 | 2.9 | | X | | |
| PDXD1 | 1.9 | | X | | X |
| PKN2 | 1.1 | | X | | |
| PLXB2 | 2.9 | X | X | | |
| PRS4 | 1.1 | | X | | |
| PRS7 | 1.0 | X | X | | |
| PSA2 | 2.0 | | X | | |
| PSA6 | 1.5 | | X | | |
| PSB1 | 1.5 | | X | | |
| PSB5 | 1.7 | | X | | |
| PSD12 | 1.5 | | X | | |
| PTN23 | 3.8 | | | | X |
| PUR6 | 1.8 | X | X | | |
| RETST | 3.2 | | X | | |
| RPB1 | 3.6 | X | | | |
| RSU1 | 1.3 | | X | | |
| SCRIB | 2.7 | X | X | | |
| SNG2 | 3.6 | | X | | X |
| SPCS3 | 1.2 | X | X | | |
| SPTN1 | 4.2 | | X | | |
| SRC8 | 1.5 | X | X | | X |
| STAM1 | 3.0 | | X | | X |
| STX4 | 2.8 | | X | | |
| STXB3 | 2.0 | | X | | |
| SYAP1 | 1.9 | | | | X |
| TANC2 | 5.9 | | X | | |
| TAP1 | 1.6 | | X | | |
| TAP2 | 1.5 | | X | | |
| TB182 | 3.0 | X | X | | |
| TERA | 1.6 | X | | | |
| TLN2 | 2.7 | | X | | |
| TMF1 | 7.7 | | | | X |
| TMOD3 | 1.6 | | X | | |
| TXLNG | 2.3 | X | | | |
| UBE4A | 4.4 | | X | | |
| UGPA | 1.6 | | X | | X |
| UN45A | 1.2 | | X | | |
| VINC | 1.9 | X | X | | |
| YTHD3 | 3.6 | X | | | |
| ZW10 | 7.5 | | X | | |

### Figure S8 (Related to Figure 5A)

#### The 141 autophagy-related hits

| Acc | Log2 fold | Reported association with LC3 | LIR | High confidence - Volcano Plot (FDR) | Le garroue et al. | Dengjel et al. | Gao et al | Mancias et al. | Autophagy Pathway/ autophagosome localisation | Biological processes |
|---|---|---|---|---|---|---|---|---|---|---|
| A16L1 | 3.0 | X | X | | | | | | X | |
| AAKG1 | 1.9 | | | | | X | X | | X | |
| ABCD1 | 1.2 | X | | | | | | | | |
| ABCE1 | 1.5 | | | X | X | X | | | | |
| ACLY | 1.8 | | | X | X | X | | | | |
| ACTN4 | 4.5 | | | | X | X | | | | X |
| AKAP1 | 7.6 | | | | | | | | | |
| ANM5 | 2.6 | | | X | | X | | | | X |
| ANXA7 | 2.1 | X | | | | X | | | | |
| ATF6A | 5.3 | | | | | | | | | |
| ATG3 | 2.4 | X | | | | | | | X | |
| ATG7 | 2.7 | X | | | | | X | | X | |
| ATG9A | 5.0 | | | | | | X | X | X | |
| BACH1 | 5.4 | | | | | | | | | X |
| BAG3 | 1.3 | | X | | | X | | X | | |
| BAG6 | 2.7 | X | X | X | | | | | | X |
| BCAR1 | 7.3 | | | X | | | | | | X |
| BIRC6 | 3.1 | X | X | X | | | | | | |
| CACO2 | 7.3 | X | | | | X | | X | X | |
| CAN1 | 2.9 | | | | | | | | | |
| CAPZB | 1.8 | | | | X | X | | | | |
| CCD50 | 2.8 | X | | | | | | | | |
| CIP2A | 2.7 | | | | | | | | | |
| CLAP2 | 2.2 | | X | | | | | | | |
| CLU | 1.5 | X | X | | X | X | | X | | |
| CNOT1 | 3.2 | | | | | X | | | | X |
| CNOT3 | 4.2 | | | | | | | | | X |
| COG4 | 7.2 | | | | | | | | | |
| CP1B1 | 3.6 | | | | | X | | | | |
| CSN8 | 3.5 | | | | | | | | | |
| DCTN1 | 1.6 | | | | X | | | | | X |
| DCTN2 | 1.9 | | | X | | | | | | X |
| DDB1 | 2.3 | | | | | X | | | | |
| DIAP1 | 1.8 | | | | | | | | | X |
| DICER | 4.3 | | | | | | | | | |
| DJC13 | 2.2 | | | | X | X | | | | |
| EEA1 | 2.9 | X | | X | | X | | | | |
| EGFR | 1.5 | X | X | | X | | | | | |
| EIF3C | 1.2 | | | | | | | | | |
| EIF3G | 1.7 | | | | X | | | | | |
| ELP3 | 1.2 | | | | | | | | | |
| ESYT1 | 2.8 | | | X | X | | | | | |
| FANCI | 1.5 | | | | | | | | | X |
| FAS | 1.1 | | | | X | X | | | | |
| FKBP8 | 1.2 | X | X | | | X | | | | |
| FLII | 2.8 | | | X | | X | | | | |
| FLNA | 1.6 | | | X | X | X | | | | |
| FND3A | 2.0 | | X | | | | | | | |
| FYCO1 | 2.1 | X | X | | | | | X | X | |
| GBF1 | 5.6 | | X | X | | X | | | | X |
| GCR | 1.7 | | X | | | | | | | X |
| GOGA2 | 2.4 | | X | | | X | | | | X |
| GOGA4 | 6.7 | | | | | | | | | X |
| GOPC | 6.0 | | | | | | | X | | |
| HECD1 | 4.2 | | | X | | | | | | |
| HERC1 | 5.6 | | X | | | | | | | |
| HGS | 3.2 | X | | | | X | | X | | |
| HPRT | 2.0 | | | | | X | | | | |
| HUWE1 | 3.8 | | | X | | X | | | | |
| ICAL | 2.2 | | | X | | | | | | |
| IF4G1 | 1.1 | | X | | X | X | | | | X |
| IMA5 | 3.7 | | | | | | | | | |
| IRAK1 | 3.3 | | | | | | | | | |
| IRS2 | 2.7 | | | X | | | | | | |
| ITPR1 | 3.4 | | | | | X | | | | |
| ITPR3 | 3.9 | | | | | X | | | | |
| KEAP1 | 5.3 | X | | | | | | X | | |
| KINH | 2.6 | | | X | X | | | | | X |
| KPCI | 3.5 | | | X | | X | | | | X |
| LRBA | 5.9 | X | X | | | | | | | |
| MACF1 | 3.1 | | X | | | X | | | | |

| Acc | Log2 fold | Reported association with LC3 | LIR | High confidence - Volcano Plot (FDR) | Le garroue et al. | Dengjel et al. | Gao et al | Mancias et al. | Autophagy Pathway/ autophagosome localisation | Biological processes |
|---|---|---|---|---|---|---|---|---|---|---|
| MALT1 | 8.3 | | | | | | | | | |
| MBOA7 | 4.2 | | | | | | | | | |
| MLP3B | 1.4 | X | | | | X | X | X | X | |
| MTOR | 3.5 | | | | | | | | X | X |
| MYH9 | 1.3 | | | | X | | | | | X |
| MYOF | 1.6 | | | X | X | | | | | |
| NCOR1 | 5.3 | X | X | | | | | | | |
| NEK9 | 4.3 | X | | X | | | | | | |
| NFKB1 | 6.0 | | | | | | | | | |
| NU214 | 2.3 | | X | | | | | | | |
| OGA | 2.8 | | | X | | | | | | |
| OGT1 | 3.0 | | | X | | | | | | |
| PA24A | 1.3 | | | | | | | | | |
| PAR12 | 3.0 | | | | | | | | | |
| PAWR | 2.3 | | | | | X | | | | X |
| PCY1A | 2.5 | | | | | | | | | |
| PDC6I | 2.5 | | | X | X | X | | X | | X |
| PI3R4 | 1.4 | X | | | | | | | X | |
| PLEC | 1.4 | | | | X | X | | | | |
| PLIN3 | 1.5 | | | X | | | | | | |
| PLIN4 | 2.8 | | | X | | | | | | |
| PLK1 | 7.6 | | | | | | | | | X |
| PTN12 | 3.6 | | | X | | | | | | |
| PSDE | 1.2 | | | | | X | | | | |
| RB3GP | 4.6 | X | | | | | | | | |
| RBGPR | 2.2 | | | X | | X | | X | | |
| RMD3 | 2.9 | | | | | | | | | |
| ROCK1 | 6.5 | | | | | X | | | | X |
| ROCK2 | 3.6 | | | | | X | | | | X |
| S38A1 | 2.6 | | | | | X | | | | |
| S38A2 | 1.3 | | | | | X | | X | | |
| SAC1 | 2.1 | | | | | | | | | |
| SC23A | 2.5 | | | | | X | | | | X |
| SC23B | 2.5 | | | | | | | | | |
| SC24A | 3.9 | | X | | | | | | | X |
| SC24B | 4.8 | | | X | | | | | | X |
| SC24C | 2.4 | | | X | X | X | | | | X |
| SC24D | 4.5 | | | | | | | | | X |
| SC31A | 2.1 | | | | X | X | | | | X |
| SEPT9 | 1.3 | | X | | | | | | | |
| SQSTM | 3.6 | X | X | | X | X | | X | X | |
| STA5A | 2.2 | | | | | | | | | |
| STAM2 | 8.2 | | | | | X | | X | | |
| STAT3 | 1.1 | | | | | | | | | |
| STK24 | 3.3 | | | | | X | | | | |
| TAB2 | 5.7 | | | | | | | | | |
| TAXB1 | 7.8 | X | X | | | X | | X | | |
| TBC15 | 2.6 | X | | | | | | X | | |
| TBCD5 | 5.1 | X | X | | | | | | X | |
| TBD2A | 6.5 | X | | | | | | | | |
| TELO2 | 3.0 | | | | | | | | | |
| TF65 | 3.1 | | X | X | | | | | | |
| TFG | 5.7 | | X | X | | X | | | | X |
| TNIP1 | 4.6 | | X | | | | | X | | |
| TPPC8 | 1.9 | | | | | | | | | |
| TRXR1 | 2.9 | | | | X | | | | | |
| TS101 | 4.2 | | | X | | X | | X | | X |
| UBA6 | 2.3 | X | | | | | | | | |
| UBE3A | 3.1 | | | | | | | | | |
| UBP19 | 9.0 | | | | | | | | | X |
| UBP24 | 5.8 | | | | | | | | | |
| UBR4 | 2.0 | X | X | X | | X | | | | |
| USP9X | 2.9 | | | X | | X | | | | X |
| VASP | 1.5 | | | | | X | | | | X |
| VP13C | 3.7 | | | | | | | X | | X |
| WBP2 | 5.6 | | | | | | | X | | |
| WDR6 | 3.7 | | | | | | | | X | |
| WDR81 | 2.5 | X | | | | | | | X | |
| WNK1 | 2.7 | | | X | | | | | | |
| ZNT1 | 2.8 | | | | | | | | | |

### Figure S9 (related to Figure 5A)

**28 hits found in the iLIR database that are related to autophagy**

| Previously reported to be related to autophagy | | | | | |
|---|---|---|---|---|---|
| Association with LC3 reported | | | No reported association with LC3 | | |
| | Log2fold | References | | Log2fold | References |
| **A16L1** | 3.0 | Fujita et al., 2008 | **BAG3** | 1.3 | Cao et al., 2017 |
| **BAG6** | 2.7 | Sebti et al., 2014 | **CLAP2** | 2.2 | Negrete-Hurtado et al., 2020 |
| **BIRC6** | 3.1 | Jia and Bonifacino, 2019 | **FND3A** | 2.0 | Manfrini et al., 2020 |
| **CLU** | 1.5 | Zhang et al., 2014 | **GBF1** | 5.6 | Naydenov et al., 2012 |
| **EGFR** | 1.5 | Tan et al., 2015 | **GCR** | 1.7 | Lee et al., 2018 |
| **FKBP8** | 1.2 | Bhujabal et al., 2017 | **GOGA2** | 2.4 | Biazik et al., 2015 |
| **FYCO1** | 2.1 | Pankiv et al., 2010 | **HERC1** | 5.6 | Schwarz et al., 2020 |
| **LRBA** | 5.9 | Martinez-Jaramillo and Trujillo-Vargas, 2020 | **IF4G1** | 1.1 | Ramírez-Valle et al., 2008 |
| **NCOR1** | 5.3 | Saito et al., 2019 | **MACF1** | 3.1 | Sohda et al., 2015 |
| **SQSTM** | 3.6 | Lippai and Lyw, 2014 | **NU214** | 2.3 | Simioni et al., 2016 |
| **TAXB1** | 7.8 | Tumbarello et al., 2015 | **SC24A** | 3.9 | Jeong et al., 2018 |
| **TBCD5** | 5.1 | Popovic et al., 2012 | **SEPT9** | 1.3 | Wang et al., 2016 |
| **UBR4** | 2.0 | Tasaki et al., 2013 | **TF65** | 3.1 | Nopparat et al., 2017 |
| | | | **TFG** | 5.7 | Steinmetz et al., 2020 |
| | | | **TNIP1** | 4.6 | Paul et al., 2012 |

### Figure S10 (Related to Figure 5A)

**141 hits previously shown to be related to autophagy**

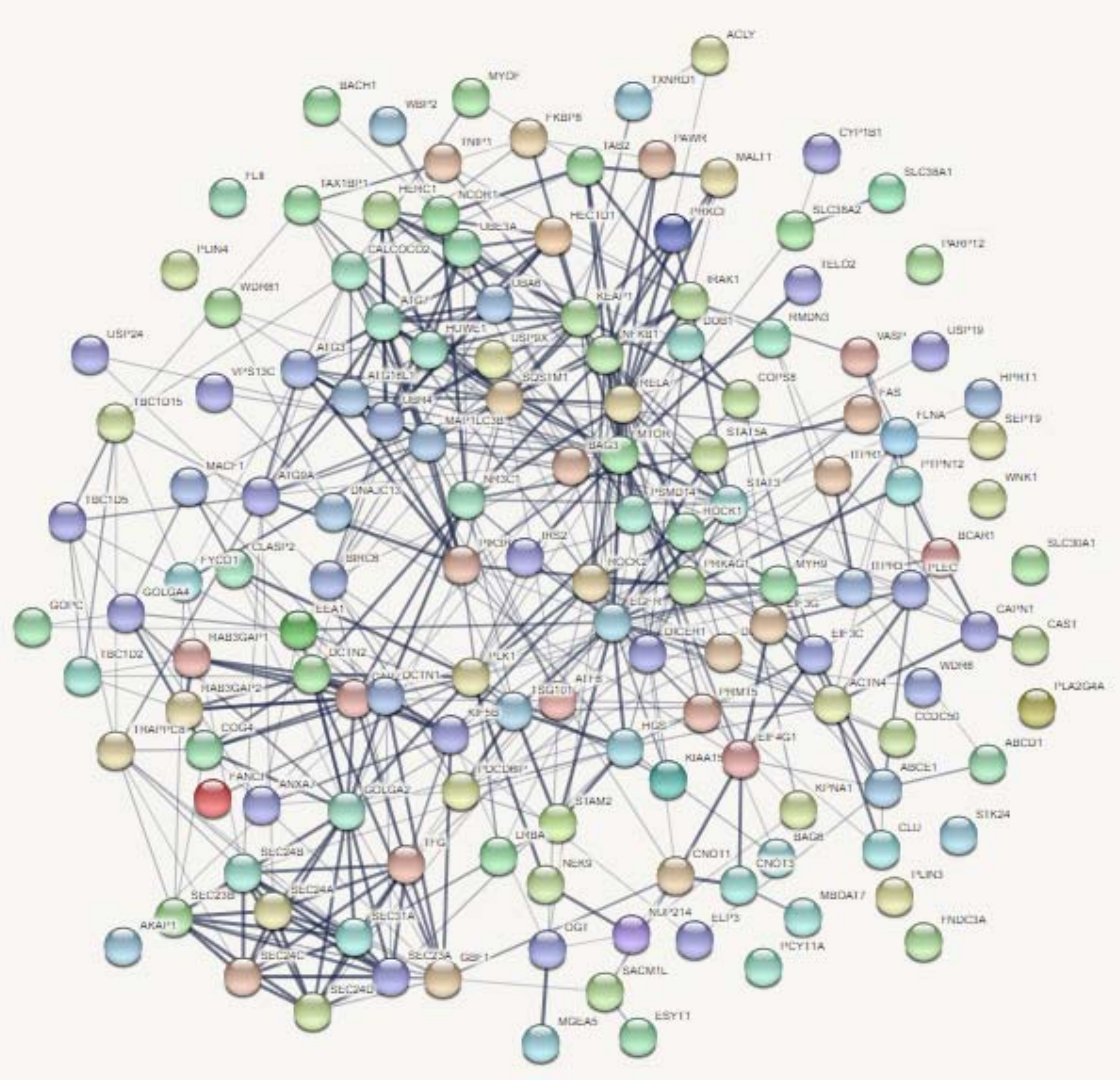

### Figure S11 (Related to Figure 6)

**42 hits not previously shown to be related to autophagy or associated with LC3 but with putative LIR domain**

| | | Trafficking | | | Cell cycle | | | RNA |
|---|---|---|---|---|---|---|---|---|
| | Log2fold | Biological process | Localisation | Pathway | Biological process | Localisation | Pathway | Pathway |
| **AASD1** | 1.7 | | | | | | | X |
| **ACAP2** | 4.2 | | X | X | | | | |
| **ANKL2** | 4.1 | | | | X | | | |
| **ANX11** | 1.6 | | X | | | X | X | |
| **BCAR3** | 7.9 | | X | | | | | |
| **BUB1B** | 2.6 | | X | | X | X | X | |
| **CNO10** | 3.3 | | X | | X | | | X |
| **CYTSB** | 6 | | | | | | | |
| **DLG1** | 5.4 | | X | | X | | | |
| **DLGP5** | 4.6 | | X | | X | X | X | |
| **DNMBP** | 5.8 | | | X | | | | |
| **EIF2D** | 2.1 | | X | | | | | X |
| **EP15R** | 2.9 | | X | X | | | | |
| **EPIPL** | 1.9 | | X | | | | | |
| **GAPD1** | 2.6 | | X | X | | | | |
| **GOGA3** | 4.3 | | X | X | | | | |
| **GOGB1** | 9.7 | | | X | | | | |
| **INF2** | 2.3 | | | | | | | |
| **JIP4** | 3.2 | | X | X | | | | |
| **KI13A** | 6 | | X | | | | X | |
| **KIF14** | 3.8 | | X | | | X | | |
| **LIMC1** | 2.5 | | | | | | | |
| **LR16A** | 1.6 | | X | | | | | |
| **MIA3** | 4.7 | X | X | | | | | |
| **OSBP1** | 3.4 | | X | | | | | |
| **PDE3A** | 3.2 | | | | X | | | |
| **PJA2** | 4.7 | | X | | | | | |
| **PRC2C** | 2.7 | | | | | | | X |
| **R3HD1** | 2.5 | | | | | | | X |
| **RABE1** | 7 | | X | X | | | | |
| **RABE2** | 6.4 | | X | X | | | | |
| **SH3G1** | 1.5 | | X | X | | | | |
| **SPD2B** | 2.7 | | | | | | | |
| **SPG20** | 2.4 | | X | X | | | | |
| **STRN** | 3.4 | | | | | | | |
| **TACC2** | 1.7 | | X | | | | | |
| **TENS3** | 3.2 | | | | | | | |
| **TENS4** | 7.4 | | | | | | | |
| **TFG** | 5.7 | X | X | X | | | | |
| **TLN2** | 2.7 | | | | | | | |
| **UBE2O** | 3.7 | | X | | | | | |
| **UBP47** | 5.1 | | X | | X | | | |

### Figure S12

**53 hits with a putative LIR domain and part of membrane trafficking**

| | Log2fold | Reported association with LC3 |
|---|---|---|
| A16L1 | 3.0 | X |
| ACAP2 | 4.2 | |
| ANKL2 | 4.1 | |
| ANX11 | 1.6 | |
| BAG6 | 2.7 | X |
| BCAR3 | 7.9 | |
| BIRC6 | 3.1 | X |
| BUB1B | 2.6 | |
| CLAP2 | 2.2 | |
| CNO10 | 3.3 | |
| DLG1 | 5.4 | |
| DLGP5 | 4.6 | |
| DNMBP | 5.8 | |
| EGFR | 1.5 | X |
| EIF2D | 2.1 | |
| EP15R | 2.9 | |
| EPIPL | 1.9 | |
| FKBP8 | 1.2 | X |
| FND3A | 2.0 | |
| FYCO1 | 2.1 | X |
| GAPD1 | 2.6 | |
| GBF1 | 5.6 | |
| GCR | 1.7 | |
| GOGA2 | 2.4 | |
| GOGA3 | 4.3 | |
| GOGB1 | 9.7 | |
| HERC1 | 5.6 | |
| IF4G1 | 1.1 | |
| JIP4 | 3.2 | |
| KI13A | 6.0 | |
| KIF14 | 3.8 | |
| LR16A | 1.6 | |
| LRBA | 5.9 | X |
| MIA3 | 4.7 | |
| NCOR1 | 5.3 | X |
| NU214 | 2.3 | |
| OSBP1 | 3.4 | |
| PJA2 | 4.7 | |
| RABE1 | 7.0 | |
| RABE2 | 6.4 | |
| SC24A | 3.9 | |
| SEPT9 | 1.3 | |
| SH3G1 | 1.5 | |
| SPG20 | 2.4 | |
| SQSTM | 3.6 | X |
| TACC2 | 1.7 | |
| TBCD5 | 5.1 | X |
| TF65 | 3.1 | |
| TFG | 5.7 | |
| TNIP1 | 4.6 | |
| UBE2O | 3.7 | |
| UBP47 | 5.1 | |
| UBR4 | 2.0 | X |
